# L-SCRaMbLE creates large-scale genome rearrangements in synthetic Sc2.0 chromosomes

**DOI:** 10.1101/2022.12.12.519280

**Authors:** Timon Alexander Lindeboom, María del Carmen Sánchez Olmos, Karina Schulz, Cedric Kilian Brinkmann, Adán Andrés Ramírez Rojas, Lena Hochrein, Daniel Schindler

## Abstract

Optimization of the metabolic flux through heterologous pathways to improve bioproduction or utilization of alternative substrates requires both fine-tuning of non-native gene expression levels and improvement of the host genome. The SCRaMbLE system incorporated into synthetic Sc2.0 yeast strains enables a rapid approach to rearrange the genome of *Saccharomyces cerevisiae* in order to create optimized chassis. Here, we show that the light-inducible Cre recombinase L-SCRaMbLE can efficiently generate diverse recombination events when applied to Sc2.0 strains containing a linear or circular synthetic chromosome III. We present an efficient and straightforward workflow for the identification of complex rearranged synthetic chromosomes from SCRaMbLEd isolates without selection pressure. The screening method is based on novel genotyping primers, the *loxPsym* tags, which indicate not only deletions but also inversions and translocations. Long-read Nanopore sequencing is used to decode the selected genotypes and shows in conjunction with flow cytometry that large-scale karyotype alterations can be a consequence of SCRaMbLE.

## Introduction

Biology is transitioning from a science of observation toward a science of engineering. A milestone to achieve this is based on the ability to sequence genomes with great ease and within rapid timeframes (Darwin Tree of Life Project, 2022; Lewin *et al*., 2018). However, complete sequence-based knowledge still does not allow one to create life from scratch (Hutchison *et al*., 2016). But seeing life from an engineering perspective, and the constant progress in improving DNA synthesis has allowed some remarkable studies in rewriting genomes (Schindler *et al*., 2018). The synthetic yeast project (Sc2.0 project) is nearing its completion and on track to create the first fully synthetic designer eukaryote (Pretorius and Boeke, 2018). Each of the chromosomes is constructed in a designated strain and subsequently consolidated to the final yeast 2.0 (Richardson *et al*., 2017). Within the design of the Sc2.0 genome many sequence alterations were incorporated, one of the most remarkable might be the implementation of a novel *Synthetic Chromosome Recombination and Modification by LoxP-mediated Evolution* (SCRaMbLE) system (Richardson *et al*., 2017). SCRaMbLE harnesses the ability of site-specific recombinases, in particular Cre recombinase in the Sc2.0 project. Symmetrical loxP sites, hereafter referred as *loxPsym*, were inserted 3 bp downstream of most non-essential genes in the Sc2.0 design as well as on other major landmarks (i.e. centromeres). Upon induction of Cre recombinase, Cre can recombine two *loxPsym* sites and thereby cause stochastic deletion, inversion, translocation and duplications of *loxPsym*-flanked DNA sequences (Fig. 1A). Thus, a single isogenic strain can be turned into a highly diverse population that can be screened for promising candidates with improved properties (Blount *et al*., 2018; Liu *et al*., 2018; Shen *et al*., 2016) (Fig. 1B). The sequence flanked by two adjacent *loxPsym* sites is referred to as a *loxPsym* unit (LU) and this term is used hereafter for the convenience of subsequent description (Luo *et al*., 2021). SCRaMbLE has been proven as a solid tool to rapidly evolve synthetic yeast strains with phenotypes of interest (Blount *et al*., 2018; Gowers *et al*., 2020; Liu *et al*., 2018) or to generate minimized genomes (Luo *et al*., 2021). In the future, SCRaMbLE may be widely applied to probe improvements of bioproduction processes in yeast which may be transferred to other industrially relevant yeast chassis. The on-demand creation of large-scale genomic rearrangements realized by SCRaMbLE is unique and may be coupled to traditional random mutagenesis and/or laboratory adaptive evolution (ALE) after promising candidates are identified (Sandberg *et al*., 2019).

**Figure 1.**
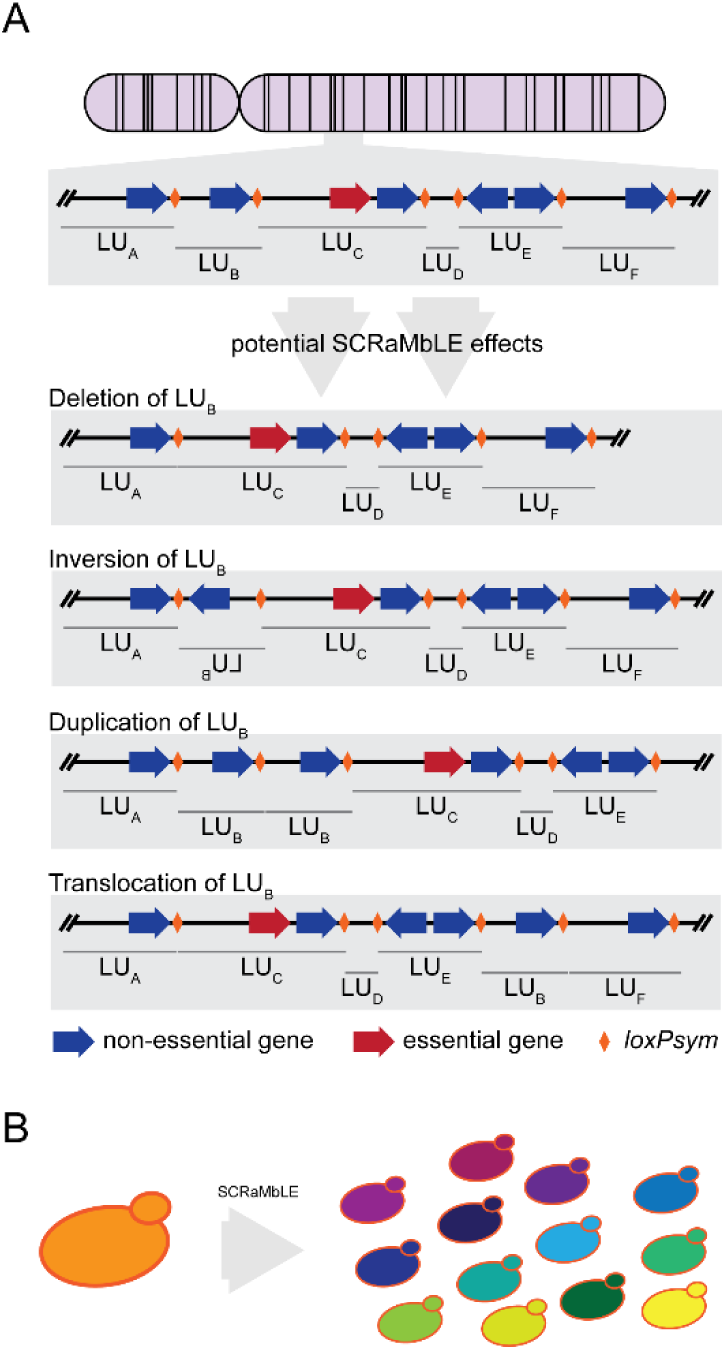
The basics of SCRaMbLE and PCRTags. **(A)** SCRaMbLE causes deletions, inversions, duplications and translocations between two *loxPsym* sites in synthetic yeast strains as depicted by LU_B_. **(B)** SCRaMbLE can turn a single isogenic synthetic yeast strain into a highly diverse population with different properties and characteristics.

However, efficient recombination of synthetic Sc2.0 chromosomes, whether for studying chromosome structure, generating minimal genomes, or optimizing chassis for biotechnological applications, imposes the following main requirements on a suitable recombination system: (1) tight control of Cre recombinase prior induction, (2) moderate Cre activity upon induction resulting in manifold and diverse recombination events, (3) reliable off-switch to generate isogenic yeast colonies post-SCRaMbLE. All these properties apply to our previously established red light-inducible Cre recombinase L-SCRaMbLE, when applied to plasmid pLM494 carrying four *loxPsym*-flanked genes of the β-carotene pathway (Hochrein *et al*., 2018). L-SCRaMbLE is based on a split Cre recombinase, whereby its N- and C-terminal halves are fused, respectively, to an N-terminal version of the photoreceptor phytochrome B (PhyBNT) and its interacting factor PIF3 from *Arabidopsis thaliana* (Hochrein *et al*., 2018) (Fig. 2A). While the recombinase is inactive in darkness or under far-red light conditions (740 nm), the split proteins will dimerize after red light induction (660 nm) and form a functional Cre, that can work on *loxPsym* recombination sites. This dimerization can be reversed via far-red light application and in this way allows a complete shut-off of Cre activity just by changing the irradiated wavelength, without manipulating growth conditions by media exchange. When previously tested on plasmid pLM494, we could show that L-SCRaMbLE achieves recombination efficiencies of up to 90%, tunable by induction time and the concentration of the chromophore phycocyanobilin (PCB) (Hochrein *et al*., 2018). The light-inducible Cre produced a large diversity of recombination events and could be completely shut off by withdrawal of the chromophore, irrespective of the light condition resulting in isogenic yeast colonies post-SCRaMbLE.

**Figure 2.**
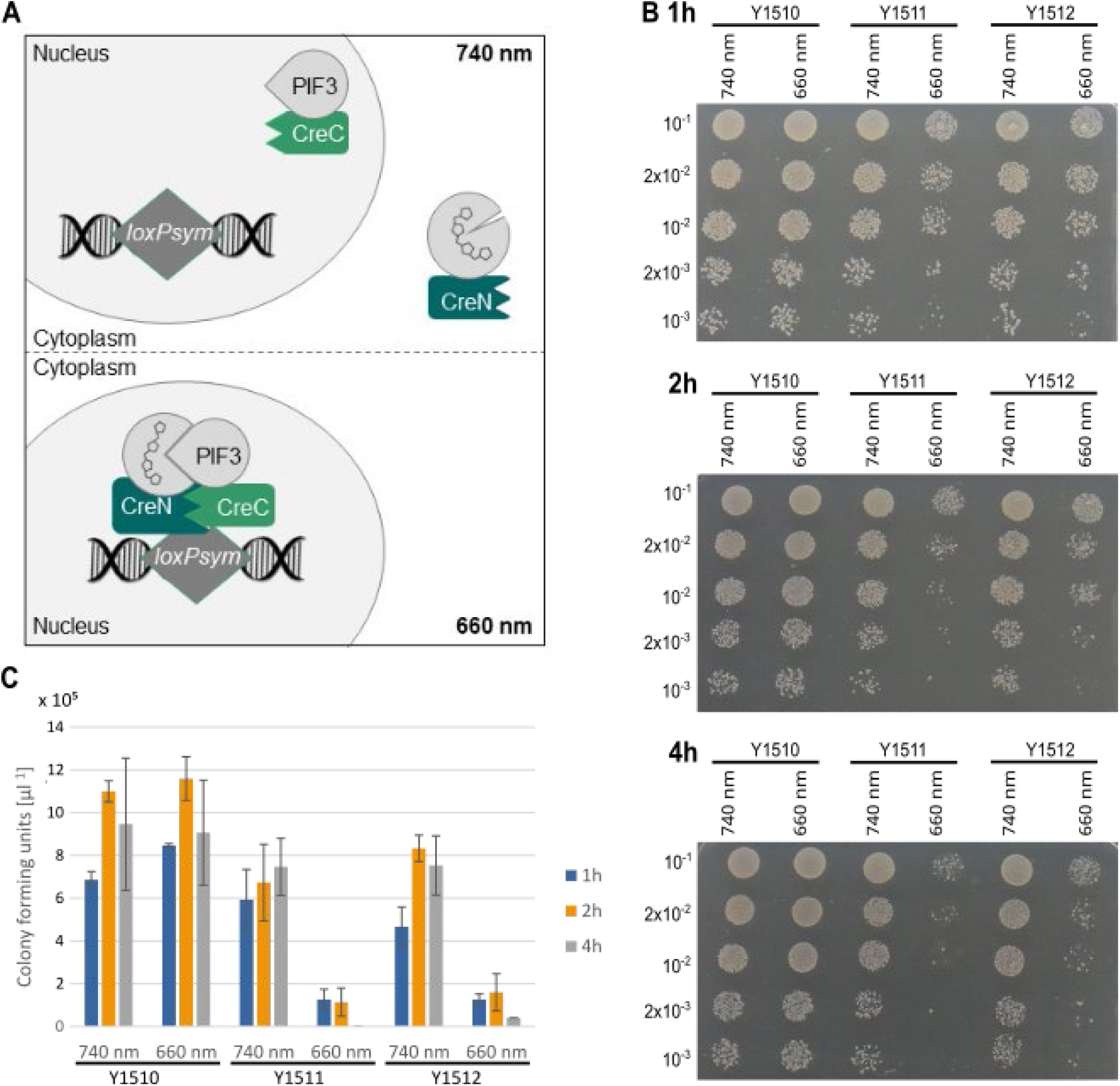
Schematic overview of L-SCRaMbLE and effect on semi-synthetic Sc2.0 yeast strains. **(A)** Mode of action of Y1510- Y1512 cells carrying the L-SCRaMbLE plasmid pLH_Scr15. L-SCRaMbLE is inactive in the dark or in far-red light conditions (740 nm) in which PhyBNT-CreN is located in the cytoplasm and PIF3-CreC is located in the nucleus. Upon induction with red light (660 nm) PIF3-CreC and PhyBNT-CreN dimerize in the cytoplasm and reconstitute a functional Cre recombinase that can translocate to the nucleus and act on *loxPsym* recombination sites. Figure reused from Fig. 1 in Hochrein *et al*., 2018. **(B)** Y1510-Y1512 cells were grown in darkness for 6 h. Induced samples were cultured in medium containing 25 µM PCB and irradiated by a 5-min red light pulse, followed by 10-sec red light pulses every 5 min for 1 h, 2 h and 4 h, respectively. Non-induced samples were cultured in darkness for 1 h, 2 h and 4 h, respectively. After the indicated time, serial dilutions were performed and 20 µl of each sample was spotted on SD-Leu plates. Each experiment was performed in three independent replicates. Pictures show one of three biological replicates. **(C)** Bar blots show the mean values of colony forming units per µl calculated from three biological replicates. Error bars indicate standard deviation.

Here, we show for the first time that the light-inducible Cre recombinase L-SCRaMbLE can tightly regulate recombination of Sc2.0 strains holding a linear or circular synthetic version of chromosome III (synIII), generating manifold SCRaMbLE events (Annaluru *et al*., 2014). Based on previous results showing that circular synthetic chromosomes exhibit more complex recombination events post-SCRaMbLE compared to linear ones (Wang *et al*., 2018), we included a circular version of synIII in our study to test whether this effect can also be observed with L-SCRaMbLE. We developed a novel workflow for the high-throughput analysis of Sc2.0 strains to identify and analyze promising candidates showing large-scale genomic rearrangements. To allow efficient pre-screening of SCRaMbLEd cultures via endpoint qPCR to identify strains showing multiple recombination events, we introduce a novel type of SCRaMbLE genotyping primers, termed loxTags. LoxTags were designed to bind upstream and downstream of *loxPsym* sites, and thus indicate three of the four possible structural variations: deletion, inversion and translocation. This is an advantage over the established Sc2.0 PCRTags, which were developed to distinguish between wild-type and synthetic DNA and only allow detection of the presence or absence of a LU (*cf*. Fig. 3A-C). To facilitate the design of custom diagnostic PCR genotyping primers for high-throughput analysis, we developed a computational tool (*qTagGer*). Pre-screened candidates showing promising recombination events were further analyzed by Nanopore sequencing, flow-cytometry and pulsed-field gel electrophoresis.

**Figure 3.**
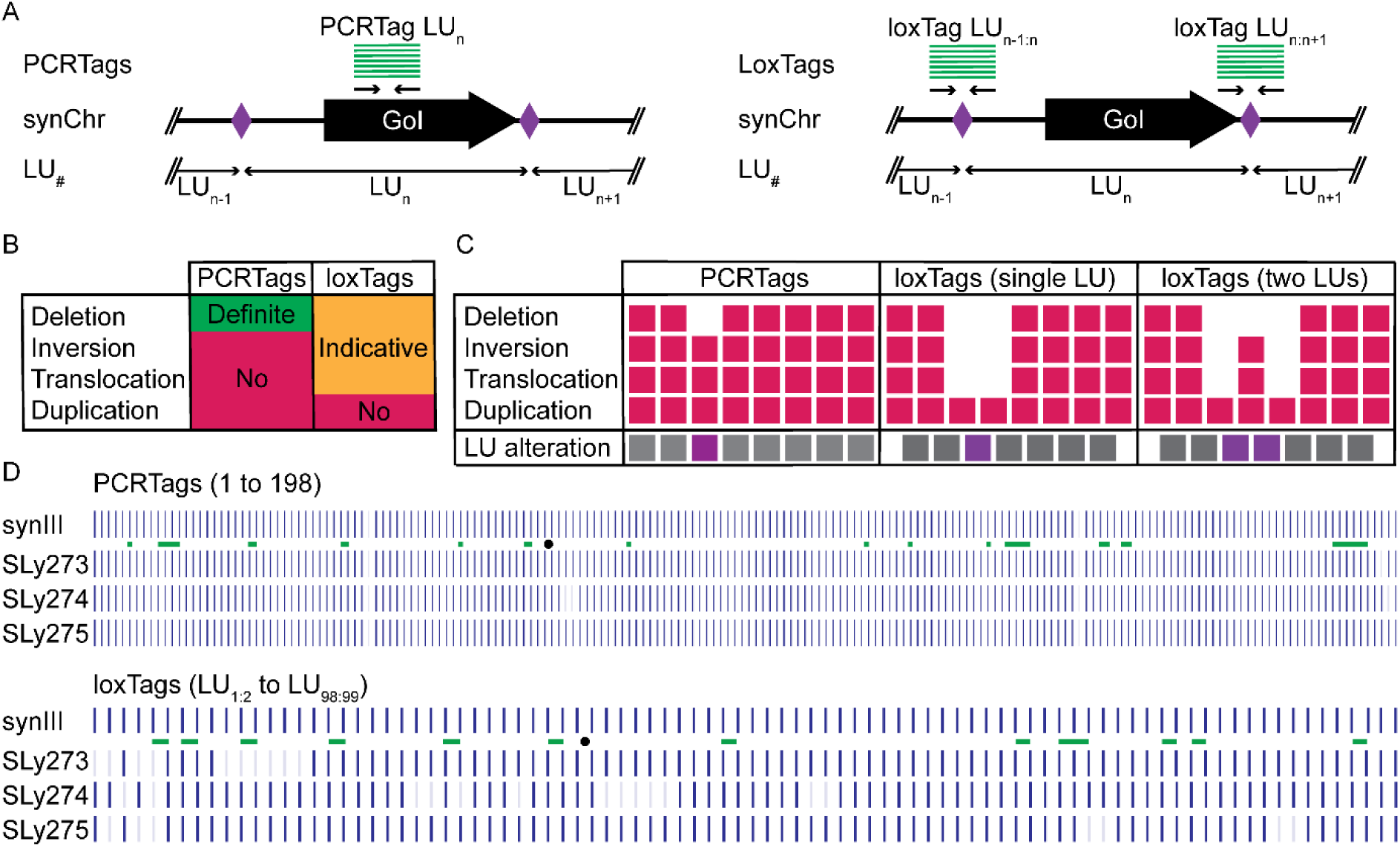
Principles, characteristics and application of PCRTags and LoxTags. **(A)** PCRTag and loxTag basic principles. PCRTags are recoded sequences within a gene to distinguish synthetic from wildtype DNA. LoxTags generate an amplicon spanning distinct *loxPsym* sites. **(B)** Comparison of PCRTag and loxTag outputs. PCRTags can only indicate the presence or absence of a tested DNA sequence. In contrast to the PCRTags loxTags can indicate a wider variety of SCRaMbLE events, but the results are only indicative and would need verification for example by Nanopore sequencing. **(C)** Theoretical outcome of PCRTag and loxTag analysis. Red boxes depict theoretical endpoint qPCR results obtained using either PCRTags or loxTags based on a reference where an LU alteration occurred (purple box). PCRTags only detect deletion of LUs while loxTags can indicate deletions, inversions and translocations. **(D)** Exemplary results of endpoint qPCR with PCRTags and loxTags. Dark blue color indicates obtained endpoint qPCR signal, absence of signal is shown by faint bar. LoxTags indicate more events compared to the PCRTags. Strikingly both datasets indicate that a larger region right to the centromere in SLy274 is lost. Black dot indicates position of the centromere, green bars indicate location of essential genes/essential LUs.

## Results

### L-SCRaMbLE generates manifold structural variations in synthetic yeast chromosomes

The red light-inducible Cre recombinase L-SCRaMbLE is built from an optical dimerizer from *A. thaliana* fused to a split version of Cre recombinase (Fig. 2A). While non-active in darkness or far-red light conditions, L-SCRaMbLE can be rapidly activated upon short red light pulses. Importantly, Cre activity can be completely switched off again by illumination with far-red light. Here, we show that L-SCRaMbLE is ideally suited to generate manifold recombination events on yeast strains holding a synthetic version of chromosome III (synIII). SynIII is 272,195 bp in size, approximately 44 kb shorter than the native chromosome, and holds 98 *loxPsym* sites (Richardson *et al*., 2017). The circular synIII created within this study is 271,793 bp, holds 98 *loxPsym* sites and shows no obvious phenotypic changes compared to the linear synIII. Since deletions of essential genes will cause lethality, growth assays were performed to show that L-SCRaMbLE can efficiently act on genome integrated *loxPsym* sites in a time-dependent manner (Fig. 2B and C). To this end, wild type BY4742, as well as semi-synthetic yeast strains holding either a linear or a circular version of synIII were transformed with L-SCRaMbLE plasmid pLH_Scr15, yielding yeast strains Y1510-Y1512, respectively. These strains were either induced with recurring red light pulses (660 nm) for 1 h, 2 h and 4 h, respectively, or kept in non-inducing conditions (740 nm) for the same period, spotted on plates in serial dilutions and analyzed for their growth on SD-Leu plates. While, as expected, no difference in colony number between induced and non-induced samples can be detected in the wild-type strain, the synIII strains show a clear reduction in cell viability already one hour after red light treatment. However, the effect becomes more pronounced with increasing induction time (Fig. 2B and C). Our results suggest that one hour of induction is sufficient to obtain a diverse pool of SCRaMbLEd yeast cells.

### Novel PCRTags expand the detection of SCRaMbLE events in synthetic yeast

The rewriting of the Sc2.0 synthetic chromosomes includes the recoding of genes to introduce PCRTags. PCRTags are short nucleotide sequences allowing the generation of specific amplicons to prove the presence of synthetic DNA (Richardson *et al*., 2017) (Fig. 3A). Initially PCRTags were used to distinguish the native from the synthetic DNA during the construction of the synthetic chromosomes. Additionally, they have proven useful for genotype screening of SCRaMbLEd candidates to explain phenotypes and identify promising strains for whole genome sequencing and subsequent genotype reconstruction (Liu *et al*., 2018; Luo *et al*., 2021; Ong *et al*., 2021). A cost and time efficient way to perform this analysis is high-throughput endpoint qPCR to obtain information about the genotype (Mitchell *et al*., 2015). We used the designed Sc2.0 primers to screen candidates after L-SCRaMbLE. Endpoint qPCR analysis allows the detection of missing PCRTags which correspond to deleted LUs during the SCRaMbLE procedure (*cf*. Fig. 3B-D). However, the current PCRTags give only information about the presence or absence of a given amplicon corresponding to a targeted gene (Fig. 3A-C). PCRTags do not detect other possible SCRaMbLE outcomes such as inversions, translocations or duplications and not every LU contains a PCRTag. We therefore designed and tested novel genotyping primers allowing us to test if the order or orientation of LUs are altered (Fig. 3A). The diagnostic primer pairs are designed to span the individual *loxPsym* sites and therefore can give information in regard to potential inversions, deletions and translocations but do not allow the detection of duplications. To distinguish the novel diagnostic amplicon primers from the Sc2.0 PCRTags we termed these primer pairs loxTags. To allow high-throughput primer design we created a computational tool for diagnostic qPCRTag generation (qTagGer). qTagGer creates a set of diagnostic PCR amplicons for a user defined motif (i.e. *loxPsym*) within a user defined input sequence and utilizes Primer3 (Rozen and Skaletsky, 2000) for primer design. qTagGer checks the designed primer pairs against the provided reference to validate their specificity (for details see material and methods and Fig. S1). We designed the loxTags for all synthetic chromosomes, including the tRNA neochromosome (Supporting Data S1) which carries *rox* sites recognized by the Dre recombinase, an orthogonal but not cross acting recombinase system (Anastassiadis *et al*., 2009; Schindler *et al*., 2022). In this way we can prove the versatility of our tool which can be used with any custom DNA sequence motif. We provided all loxTags for the Sc2.0 chromosomes and the roxTags for the tRNA neochromosome in the supplementary data as resources for the Sc2.0 community (Supporting Data S1). As a proof of concept, we ordered and validated the synIII loxTags (Fig. 3D and Fig. S1). Single LU inversions, deletions or translocations would be indicated by at least two consecutive missing amplicons, which is indeed what we observed in our initial set of tested L-SCRaMbLE candidates (Fig. 3C, D). However, larger structural variations may have a different pattern of multiple loxTags missing (deletion) or only two distant loxTags missing (large inversions or translocations). Comparing the loxTags with the PCRTags we can detect more potential structural variations and confirm the loss of a larger region right of the centromere in SLy274 (Fig. 3D). The results show that loxTags have an advantage in detecting structural variations in SCRaMbLEd candidates. Using the loxTags for an initial screen of candidates increases the throughput, lowers the cost and may be sufficient to identify candidates for subsequent whole genome sequencing.

### L-SCRaMbLE activity on the linear synthetic chromosome III

The synIII strain contains the ura3Δ0 allele and a non-functional *URA3* at the HO locus (Brachmann *et al*., 1998; Mitchell *et al*., 2017). Both alleles were removed in two consecutive rounds of CRISPR/Cas9 genome editing resulting in the intermediate SLy053 and final strain SLy066. The reason for the removal is to reduce the risk of unwanted homologous recombination in downstream applications. Notably a *URA3* deletion has been described to cause sensitivity to acetic acid (Ding *et al*., 2013). The L-SCRaMbLE plasmid (pLH_Scr15) was transformed into SLy066 resulting in strain Y1511. Based on our L-SCRaMbLE activity assay results (*cf*. Fig. 2B, C), we induced L-SCRaMbLE for one hour before plating dilution series to isolate single colonies. Twelve candidates were isolated and subsequently subjected to loxTag analysis (Fig. 4A). The loxTags indicate a broad variation of events indicating all possible SCRaMbLE outcomes, except for duplications due to technical limitation. To determine the SCRaMbLE events by Nanopore sequencing we selected the eight strains with the highest diversity detected by loxTag genotyping (SLy241-248). We purposefully selected for highly diverse L-SCRaMbLE activity and did not consider any phenotypic data. Nanopore sequencing resulted on average in a 42-fold coverage for the linear synIII strains (Table S1). *De novo* assembly using canu (Koren *et al*., 2017) and manual curation resulted in a single synIII derivative for each strain. To visualize the structural variations, we used SCRaMbLEgrams (Shen *et al*., 2016) where each color of the triangles corresponds to one LU (Fig. 4B). The supporting information further provides dot plots as an alternative visualization strategy (Fig. S2). Both approaches indicate alterations to the reference and visualize deletion, inversion, duplication and translocations respectively, except for strain SLy246 which seems to have no alterations. To validate our assemblies we used pulsed-field gel electrophoresis (PFGE) to confirm the obtained assembly sizes of the synIII derivatives (Fig. 4C and Fig. S3). Because synIII (272.9 kb unSCRaMbLEd) is too close in size to chromosome VI (270.2 kb) to distinguish between the two, we used a circular synIII derivative (SLy117, details next section) as control in the PFGE. Circular chromosomes do not migrate in the gel and therefore the control allows us to distinguish between a single band from a double band. The size of our assembled SCRaMbLEd synIII chromosomes matches the observed sizes in the PFGE.

**Figure 4.**
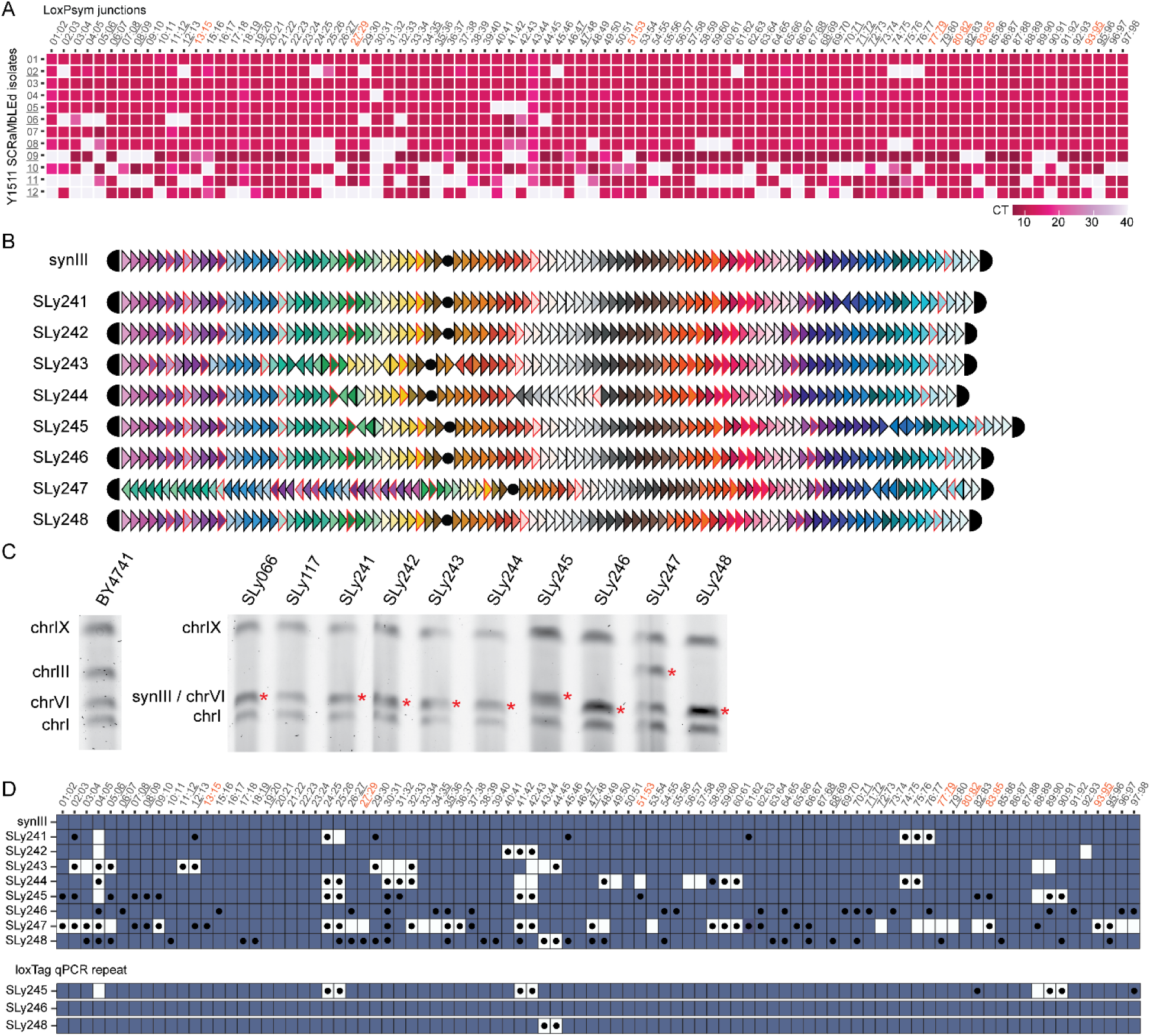
L-SCRaMbLE causes structural variations on chromosome level. **(A)** Results of loxTag analysis of tested candidates. Gray boxes indicate no crossing of the cycle threshold indicating the absence of the respective specific amplicon. The eight underlined candidates are selected for Nanopore sequencing based on a high number of observed variations, clear differences between the selected candidates, and are further on referred to as SLy241 to SLy248. Orange *loxPsym* junctions indicate loxTags which span 2 *loxPsym*, essential LUs are indicated by underlined numbers. **(B)** Visualization of synIII SCRaMbLE alterations, each colored triangle represents one distinct LU. Triangles in SCRaMbLEgrams visualize amplification, inversion and translocation; the absence of a colored triangle in comparison to the parental strain indicates a deletion. Red outlined triangles mark essential LUs and the black dot the centromere LU. **(C)** Pulsed-field gel electrophoresis of tested candidates for synIII size validation. SLy245 and SLy247 show the expected size increase. SLy117 contains a circular synIII and serves as a control to distinguish a single chrVI band from the synIII/chrVI double band. **(D)** Comparison of *in silico* endpoint qPCR of *de novo* assembled synIII derivatives with the obtained results from the qPCR. Black dots show absence of loxTag in the qPCR and empty boxes indicate absence in the *in silico* data. Strains SLy245, SLy246 and SLy248 show a high amount of false positive detection of loxTag absence while in the other cases qPCR data and *in silico* data matches well. However, in case of repetition of the qPCR the *in silico* and *in vitro* results match almost perfectly.

Notably, we are aware that the qPCR method for the selection of candidates can be error prone as shown by strain SLy246 which indicated by qPCR a highly SCRaMbLEd genotype which could not be confirmed by Nanopore sequencing. Therefore, we generated the custom python script ‘visLoxP’ simulating *in silico* amplicons to visualize loxTag patterns of our assembled genomes (Fig. 4D). The loxTags were only benchmarked for the presence of amplicons in the parental synIII strain because no known altered strains were available. Besides the strains SLy245, SLy246 and SLy248 the rate for false positives (no amplicon, but amplicon expected) is relatively low, indicating a technical issue in these cases. The rate of false negatives (amplicon, but no amplicon expected) seems to have a higher likelihood. To test our assumption, we repeated the qPCR of SLy245, SLy246 and SLy248. Our analysis shows that *in silico* and *in vitro* analysis matches well in the technical repetition of the experiment. However, in cases where SCRaMbLE caused highly complex structural variations, in particular SLy247, we may not have the synIII derivative assembled correctly which could result in false positive *in silico* data.

### Circular synthetic chromosomes show increased L-SCRaMbLE complexity

We circularized the linear synIII of strain SLy066 using CRISPR/Cas9 guided homology directed repair by providing a repair template and two gRNAs cutting in the subtelomeric repeats of the left and right arm respectively resulting in strain SLy117. The circularity of synIII was verified using PCR, Nanopore sequencing and PFGE (*cf*. Fig. 4C). The L-SCRaMbLE plasmid (pLH_Scr15) was transformed into SLy117 resulting in strain Y1512. L-SCRaMbLE with Y1512 was performed in parallel to Y1511 to guarantee identical settings. Twenty-four candidates were isolated and subjected to loxTag analysis (Fig. 5A). Eleven candidates (strains SLy249-259) of Y1512 were selected based on endpoint qPCR genotyping results in the same manner as performed for Y1511 candidates and subjected to the same Nanopore sequencing experiment. The genome coverage for the circular synIII strains was on average 29-fold (Table S1). While all tests performed for the analysis of the linear synIII derivatives support the correctness of our predicted sequence assemblies, the circular chromosomes show highly complex structural variation which may not be solved properly with the available long read *de novo* assemblers. While canu (Koren *et al*., 2017) was sufficient to assemble the linear chromosomes, canu was previously reported to have poor performance when assembling circular chromosomes (Wick and Holt, 2019). Our attempts using canu to assemble the circular synIII strains were not successful. Therefore, we used Flye (Kolmogorov *et al*., 2019) which, in case of the circular synIII derivatives, provided the most likely solutions for the SCRaMbLEd synIII chromosomes after manual curation. Flye has previously been shown to perform well for circular chromosomes (Wick and Holt, 2019). The two strains SLy252 and SLy254 show highly complex rearranged synIII chromosomes and all our assembly attempts for these strains are most likely incorrect, despite the general coverage for the two samples being >15-fold which should be sufficient to perform whole genome assembly with long-read sequencing data. We observed, and it has previously been reported, that circular chromosomes show a lower abundance in whole genome sequencing experiments which may limit our assembly attempts as well (Shen *et al*., 2022). For the aforementioned reasons we do not provide synIII assemblies for those two strains. Our data highlight that dedicated software tools to decode complex SCRaMbLEd synthetic chromosomes are necessary. Nonetheless, based on the results we obtained we can confirm that circular synthetic chromosomes show a higher degree of structural rearrangements. Notably, SCRaMbLE of the linear and circular synIII strains presented in this study were performed in parallel, allowing for comparison of the SCRaMbLE events to a certain extent. All assembled circular synIII strains show SCRaMbLE events. SLy250 and SLy259 show only moderate alterations while the other strains are highly rearranged showing all types of SCRaMbLE events, which becomes particularly evident in the SCRaMbLEgrams and dot plots (Fig. 5B and S4). Our data show clearly that for circular chromosomes the number and complexity of SCRaMbLE events is higher compared to the linear chromosomes. Interestingly, in the case of the circular synIII assemblies, the *in silico*data matches the *in vitro* data well (Fig. 5C). False positives in the *in vitro* data are almost not present, but on the contrary it seems that visLoxP indicates no amplicons for many LUs which either indicate false negatives in the qPCR or potential errors in the *de novo* assembly of the SCRaMbLEd chromosomes.

**Figure 5.**
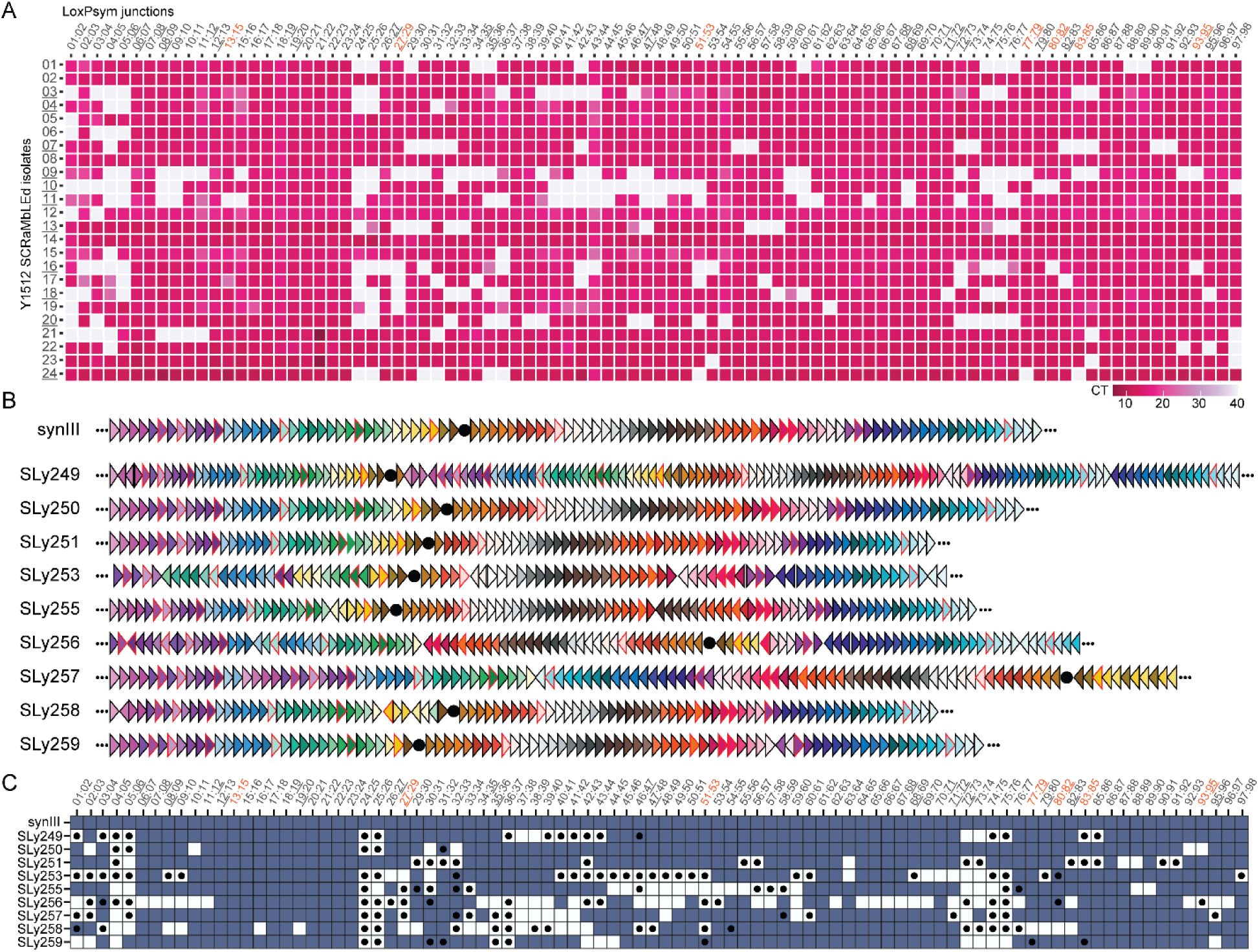
Circular chromosomes show a higher degree of L-SCRaMbLE activity. **(A)** Results of loxTag analysis of tested candidates. The eleven underlined candidates are used for Nanopore sequencing based on a high number of observed variations and clear differences between the selected candidates. The candidates are further on referred to as SLy249 to SLy258. Orange *loxPsym* junction PCRs indicate loxTags which span 2 *loxPsym*, essential LUs are indicated by underlined numbers. **(B)** Visualization of synIII SCRaMbLE alterations. Colored triangles in the SCRaMbLEgrams visualize amplification, inversion and translocation. Absence of a color in comparison to the parental strain indicates deletions. Red outlined triangles indicate essential LUs and the black dot the centromere position. Dotted lines indicate the circular nature of the chromosome. **(C)** Comparison of *in silico* endpoint qPCR of assembled synIII derivatives with the obtained results from the qPCR. Black dots show absence of loxTag in the qPCR and empty boxes indicate absence in the *in silico* endpoint qPCR. In comparison to the PCR for the linear synIII derivatives false positives detection of loxTag is relatively low. The *in silico* PCR suggests that there is a higher number of false negatives, which could also be caused by incorrect assemblies of the highly rearranged synthetic chromosomes. Generally, the qPCR data matches with the *in silico* data for the cases where amplicons are generated.

### SCRaMbLE can have a global impact on yeasts karyotype

When we analyzed the sequencing data beyond the assembly of synIII derivatives, we found an increased copy number for individual chromosomes in three instances. The abundance of reads for these chromosomes is approximately doubling in comparison to the other chromosomes indicating +1 aneuploidies (Fig. 6A). SLy245, SLy255 and SLy257 contain aneuploidies of chromosome IX, XIII and XIII respectively. Further, Nanopore sequencing results of SLy251 show a segmental duplication of roughly half of chromosome VIII. Based on this observation we decided to investigate the ploidy of all strains, as whole genome duplication has previously been described as a common stress response in yeast (Harari *et al*., 2018). We performed flow cytometry of exponentially growing cells in comparison to known haploid and diploid reference strains (BY4741, BY4743) (Fig. 6B and Fig. S5). By doing so, we discovered whole genome duplication for isolates SLy242, SLy246 and SLy248.

**Figure 6.**
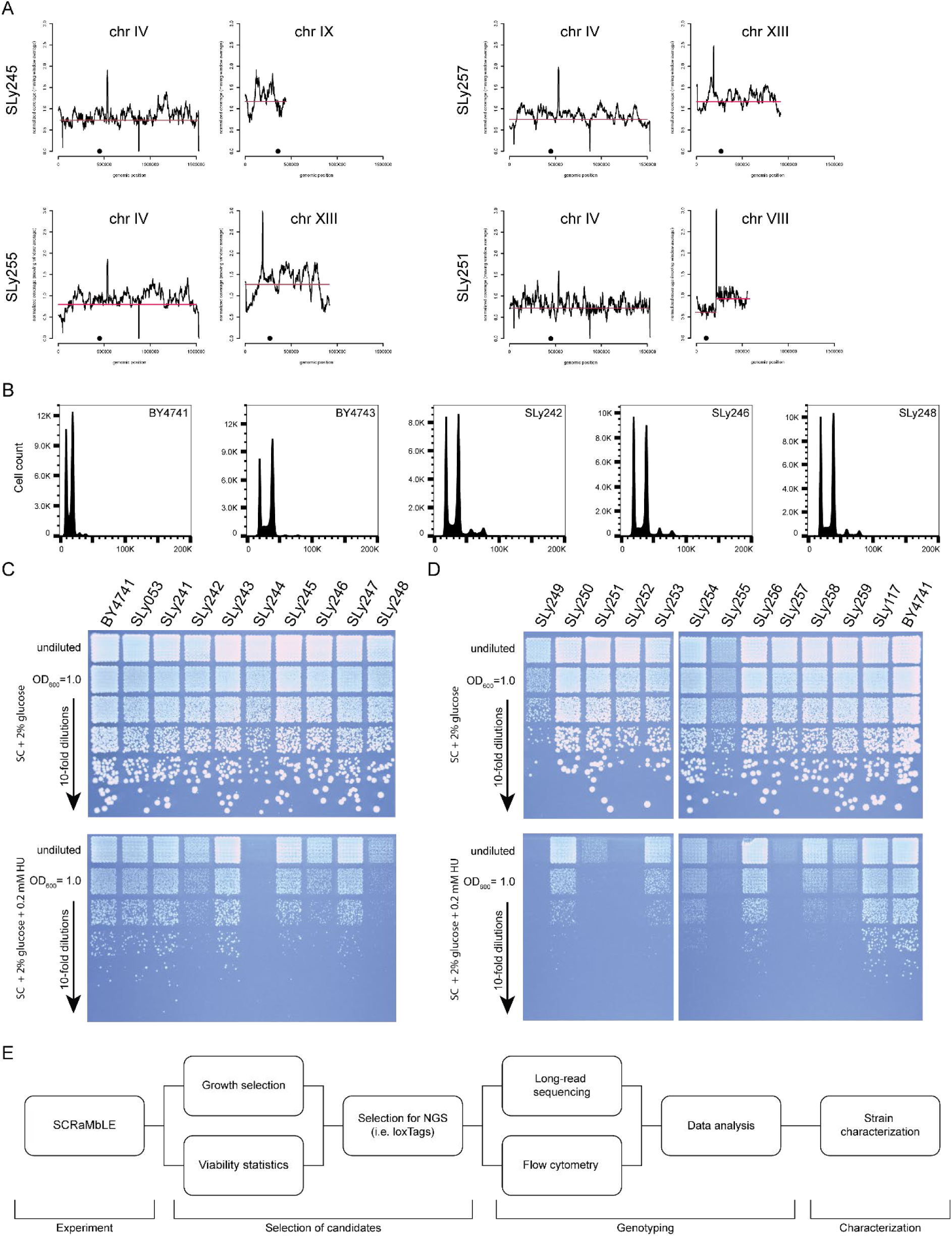
Consequences of SCRaMbLE can be complex karyotype alterations. **(A)** Aneuploidies and segmental chromosome amplification indicated by Nanopore sequencing. Red line visualizes coverage shift. **(B)** Flow cytometry identifies SLy242, SLy246 and SLy248 to have the same DNA content as BY4743. No aneuploidies can be detected for these strains in the Nanopore sequencing data indicating a whole genome duplication. **(C)** Growth assay of linear SCRaMbLE isolates on SC media and SC + 0.2 mM hydroxyurea. BY4741 and SLy053 do not show any growth differences. The whole genome duplicated strains SLy242, SLy246 and SLy248 show a decreased growth rate, consistent with previous results. The observed growth defect for SLy244 remains elusive, but the strain shows already smaller colony sizes on SC media. **(D)** Growth assay of linear SCRaMbLE isolates on SC media and SC + 0.2 mM hydroxyurea. BY4741 and SLy117 do not show any growth differences. SLy249 and SLy255 already show growth reduction on SC media. The candidates SLy255 and SLy257 have a chrXII aneuploidy and the reduced growth on hydroxyurea media is expected. Further, the strains SLy249, 251, 252 and 259 show reduced growth which cannot be pinpointed to a common cause. **(E)** Best practice workflow to perform and analyze SCRaMbLE experiments.

Importantly we have not observed any integration or recombination between synIII and the native chromosomes proving the targeted action of Cre recombinase. However, our results show that SCRaMbLE can have a variety of impacts on yeast karyotype, ranging from segmental to whole chromosome and genome duplications. It was previously reported that aneuploidies and whole genome duplication may have an impact on yeast cells in the presence of hydroxyurea (Schindler *et al*., 2022; Shen *et al*., 2022). Therefore, we performed growth assays in comparison to the parental strain on solid synthetic complete medium (SC) for the linear and circular strains (Fig. 6C and 6D). Strikingly the strains SLy242, SLy246 and SLy248 showed the expected growth defect. The same was observed for the strains SLy255 and SLy257 containing a chromosome XIII +1 aneuploidy, however, this was not the case for SLy245 with a duplication of chromosome VIII. Besides the aforementioned strains, SLy244, SLy249, SLy251, SLy252 and SLy259 show different intensities of growth defects which cannot be explained by a singular cause.

Based on our results we suggest a standard workflow to characterize SCRaMbLEd yeast isolates (Fig. 6E). The workflow starts with the SCRaMbLE experiment, including viability statistics to check for Cre activity, and the subsequent selection of isolates for in depth analysis. LoxTags help to identify relevant candidates which should be subsequently analyzed by long-read sequencing strategies to decode the genotype including aneuploidies. In parallel DNA content measurements with flow cytometry can identify whole genome duplication. Subsequently the isolated candidates can be characterized in depth in regard to their properties. Comparative growth on solid and/or in liquid medium can give insights into the growth fitness of the isolated candidates. Pulsed-field gel electrophoresis can be supportive in case of linear chromosomes to verify alterations on chromosomal scale. Such a standardized workflow becomes particularly relevant if no phenotypic screening is feasible.

## Discussion

Here, we showed for the first time that L-SCRaMbLE is a tightly regulated tool capable of causing all types of structural variations in Sc2.0 strains holding a linear or circular version of synIII. By analyzing only a small number of SCRaMbLEd strains, we were able to identify several candidates exhibiting multiple and diverse recombination events. Remarkably, without any selection for Cre activity like e.g. the ReSCuES system, which harnesses an auxotrophy marker switch to select for strains where Cre has been active (Luo *et al*., 2018). This proves that L-SCRaMbLE is an excellent tool to tightly regulate Cre activity not only at the plasmid level but also at the genome level. Further, we provide evidence that an induction of one hour is sufficient to create a highly diverse yeast population.

To allow straightforward identification and global characterization of promising candidates out of SCRaMbLEd cultures, we present an efficient high-throughput analysis workflow. This workflow uses a novel type of genotyping primers for improved screening of SCRaMbLEd yeast isolates. These loxTags detect a broader range of SCRaMbLE events compared to the classic PCRTags which only indicate deletions. Thus, the novel loxTags provide an improved decision-making tool to identify suitable candidates for subsequent characterization. LoxTags work economically and time efficiently in particular if the reaction is downscaled by laboratory automation. However, a drawback for the initial screen was the fact that we did not have known SCRaMbLEd strains at the beginning of our study to benchmark the loxTags. Now, with the first decoded genomes on hand we can benchmark the qPCR settings. By further analyzing SCRaMbLEd genomes in the future, we will be able to cover more and more LU variations and thus improve qPCR analysis of SCRaMbLEd strains. It is worth mentioning that a screening method selecting for the absence of a PCR amplicon is not ideal, but on this global scale it is the only strategy to reduce the set of candidates for Nanopore sequencing, a costly and time-consuming process in comparison to qPCR. With our approach we can screen four synIII candidates on a 384-well qPCR run lowering the costs to less than 5 € per strain (including DNA extraction etc.) and obtaining results within 90 minutes (excluding DNA extraction). Therefore, we are sure that the novel loxTags will be a helpful resource for the community. Moreover, the newly established tool qTagGer, which generates genotyping primers, may be broadly applied in other aspects where rearrangements at specific nucleotide sequences need to be studied.

In this study, long-read Nanopore sequencing has once more proven its ability to solve highly complex rearranged genomes and seems to be the gold standard to decode the complex genotypes of SCRaMbLE isolates. We have already sequenced up to 24 yeast strains on one MinION flow cell (Luo *et al*., 2021; Ong *et al*., 2021; Schindler *et al*., 2022). Our study confirms previous studies (Shen *et al*., 2022) which have shown that circular synthetic chromosomes show a much higher number of Cre-mediated recombination events than linear ones. The available standard *de novo* assembly tools used to assemble the sequences of highly complex rearranged circular chromosomes need intense manual curation and may not result in the correct solution. Novel computational analysis pipelines are necessary to improve analysis of SCRaMbLEd synthetic yeasts, in particular if the complexity of strains with multiple synthetic chromosomes increases.

Taken together, we show here that L-SCRaMbLE is ideally suited to rearrange synthetic Sc2.0 strains and thus generate a variety of different strains from single isogenic yeast strains. To identify, genotype and characterize promising candidates, we have established an efficient three-step workflow. This workflow will be important for future studies where SCRaMbLE will be employed to gain knowledge from synthetic Sc2.0 yeast strains to transfer it to non-synthetic industrially relevant *S. cerevisiae* strains. To our surprise, we discovered segmental chromosome duplication, aneuploidies and whole genome duplication highlighting the importance of performing a careful characterization of SCRaMbLE isolates to be able to draw conclusion in regard to phenotypes and for their potential transfer.

## Material and Methods

### Strains and growth conditions used in this study

All yeast strains were derivatives of *S288C* and grown at 30 °C unless otherwise specified. A full list of yeast strains is provided in Table S2. The standard *E. coli* TOP10 strain (Invitrogen) was used for propagation and archiving of plasmid DNA. Yeast cells were grown in either YEP/YPD media (10 g/L yeast extract, 20 g/L peptone and with or without 0.64 g/L tryptophan) or Synthetic Complete (SC) media (1.7 g/L yeast nitrogen base without amino acids and without ammonium sulfate, 5 g/L ammonium sulfate, 1.47 g/L SC. tripe drop-out -his, -leu, -ura; all components used are from Formedium) lacking the indicated amino acids, both supplemented with 2% glucose if not stated otherwise. Standard solid media contained 2% agar and cultures were incubated at 30 °C. Bacterial cultures were grown in LB medium at 37 °C with 100 μg/mL ampicillin. Bacterial and yeast liquid cultures were cultivated according to standard procedures in glass tubes on a custom-built roller drum (similar to New Brunswick TC-7) or on shaking incubators (Innova 42R, New Brunswick) at 200 rpm.

### Plasmids and oligonucleotides used in this study

All plasmids used in this study are listed in Table S3 and the plasmid files of the created plasmids are provided as Genbank files (Supporting Data S2). Oligonucleotides were ordered and synthesized by Integrated DNA Technologies in 25 or 100 nM scale as standard desalted oligonucleotides (Table S4). PCRTag primers were synthesized according to (Annaluru *et al*., 2014). LoxTags are listed separately in the supporting information; Table S5 for the physical synIII loxTags and Supporting Data S1 contains all predicted sequences.

### DNA purification and plasmid extraction

Plasmid DNA and PCR product purification was performed as previously according to open source magnetic bead purification procedures (Köbel *et al*., 2022; Oberacker *et al*., 2019). Yeast DNA isolation was performed using a modified Cetyl Trimethyl Ammonium Bromide extraction method applying glass beads (Fraczek *et al*., 2013) and a FastPrep96^TM^ bead beater for cell rupture. Briefly, cells were cultivated overnight in respective media, 1 mL culture were harvested at 13,400 *g* for 1 min in 2-mL screw cap tubes and resuspended in 1 mL of extraction buffer (2% CTAB, 100 mM Tris, 1.4 M NaCl and 10 mM EDTA, pH 8.0) and 425–600 mm glass beads were added approx. to 0.3 mL scale bar. Subsequently tubes processed either by vortexing at maximum speed for 15 min with the Vortex Genie 2 (Scientific Industries) with a 24 multi-tube holder adapter or with by placing the tubes into a FastPrep96^TM^ (MP Biomedicals) with custom-made adapters to process up to 2 x 48 tubes simultaneously. Cell rupture was performed with the following settings: 1,400 rpm for 15 min. Screw cap tubes were used to omit potential cross contamination caused by cell rupture in 96-well deep-well plates considering the sensitivity of qPCR assays. Tubes were then incubated for 10 min at 65 °C. Vortexing and incubation was then repeated once in order to improve yield. After the second incubation, tubes were centrifuged at 13,400 *g* for 1 min. The supernatant was transferred to 2-mL tubes containing 16 μL of 25 mg/mL RNaseA and incubated at 37 °C for 15 min. Afterwards, 700 μL of chloroform was added, the tube was mixed by inversion and centrifuged at 13,400 *g* for 2 min. The aqueous phase (∼700 µL) was transferred to a 1.5-mL reaction tube and 0.6 volumes of isopropanol was added. The tube was mixed by inversion and centrifuged as before. Isopropanol was discarded and the pellet was washed in 500 μL of 70% ethanol and centrifuged for 2 min at 13,400 *g*. Supernatant was discarded, the tubes pulse-centrifuged and residual alcohol was removed using pipette tips and pellet was dried with an open lid for 2-5 min at room temperature. The pellet was resuspended in 50 μL dH_2_O and stored at -20 °C until further use.

### Yeast transformation

Yeast strains were transformed using a modified protocol described by Gietz and Woods (2002). Briefly, yeast strains were inoculated into 10 mL of liquid media and incubated with rotation at 30 °C overnight. Overnight cultures were then re-inoculated to OD_600_ = 0.1 and incubated with rotation to a target OD_600_ of between 0.5 and 1. Cells were then centrifuged at 2,103 *g* for 5 min, washed with 10 mL dH_2_O and centrifuged again. Cell pellets were washed with 10 mL of 0.1 M LiOAc, centrifuged at 2,103 *g* for 5 min. Fifty μL of cells were mixed with 5 to 10 μL of transforming DNA, 36 μL of 1M LiOAc, 19 μL dH_2_O, 25 μL 10 mg/mL herring sperm carrier DNA and 240 μL of 44% PEG 3350. Transformations were incubated at 30 °C for 30 min before adding 36 μL of DMSO and heat-shock at 42 °C for 15 min. Cells were pelleted and incubated in 400 μL of 5 mM CaCl_2_ at room temperature for 10 min and plated onto selective media.

### Construction of gRNA plasmids and repair templates

Preparation of gRNA plasmids was done according to the protocol provided by the Ellis Lab https://benchling.com/pub/ellis-crispr-tools (Awan *et al*., 2017). Briefly, generation of gRNA plasmids was accomplished by insertion of annealed oligos into the destination plasmid pWS082 via Golden Gate cloning. Guide RNAs were designed using the online tool CHOPCHOP (Labun *et al*., 2019). Repair templates were generated either by using overlap extension PCR or subcloning of multiple fragments into pSL0014 via Golden Gate cloning and subsequent amplification with oligos SLo0101 and SLo0138. Overlap extension PCR was performed as following: 5’ and 3’ DNA segments flanking the area to be edited were amplified from genomic DNA using the corresponding designed oligonucleotides (*cf*. Table S4). The resulting fragments contain an overlap with matching T_m_ for the 5’ amplicon forward and 3’ amplicon reverse primer to fuse both fragments in a subsequent PCR. One µL of the 5’ and 3’ amplicon were used in the second PCR as template without purification resulting in the repair template amplified by the respective 5’ forward and 3’ reverse primers. The repair templates for CRISPR/Cas9 contained at least 300 bp for homologous recombination in yeast. Prior to yeast transformation, the gRNA plasmid(s) were linearized for one hour by digestion with EcoRV and repair template was generated using PCR.

### Construction of synIII strain derivatives

SynIII contains the ura3Δ0 allele and a remaining *URA3* marker at the HO locus (Brachmann *et al*., 1998; Mitchell *et al*., 2017). Both *URA3* alleles was removed using CRISPR/Cas9 in conjunction with one or three gRNAs targeting the ura3Δ0 allele (pSL0370) and *URA3* gene (pSL0096-98) respectively, supplied donor DNA serving as repair template resulted in strains SLy053 and subsequently SLy066. SLy066 was subsequently used for synIII ring chromosome formation using CRISPR/cas9 in conjunction with 2 gRNAs (pSL0271-272) targeting the subtelomeric repeats and a donor DNA serving as repair template (PCR amplified from pSL0270) resulting in strain SLy117 with a circular synIII. CRISPR/Cas9 editing in yeast was performed following the protocol outlined at https://benchling.com/pub/ellis-crispr-tools (Awan *et al*., 2017). Briefly, linear fragments encoding Cas9, gRNA(s) and repair template(s) were co-transformed into yeast strains using the aforementioned yeast transformation method. Plasmids containing gRNAs, DNA sequences used as repair template and circular synIII are provided as Genbank files in the supporting data (Supporting Data S2).

### Light-induced recombination experiments

For recombination experiments, yeast strains Y1510-Y1512 were inoculated in 2 mL SD-Leu in culture tubes and incubated shaking for 24 h (30 °C, 220 rpm). From this preculture, main cultures were inoculated to an OD600 ≈ 0.1 in 500 µL fresh SD-Leu medium in 24-well plates with transparent bottom (product no. 303008, Porvair Science Ltd, Norfolk, UK) covered with sterile aluminium sealing foil. Induced samples contain 25 µM PCB (phycocyanobilin, SC-1800, SiChem GmbH, Bremen, Germany). All samples were inactivated by a 1-min far-red light pulse (740 nm, 69 W/m^2^) and afterwards grown for 6 h in darkness (30 °C, 220 rpm). After that, non-induced samples were further maintained in darkness for 1, 2 or 4 h.

Induced samples were irradiated with a 5-min red light pulse (660 nm, 28 W/m^2^) and afterwards grown for 1, 2 or 4 h with 10-s red light pulses applied every 5 min. Light-induction experiments were conducted in a custom-made Light Plate Apparatus device (LPA) (Gerhardt *et al*., 2016). LPAs were equipped with 660 nm LEDs (product no. L2-0-R5TH50-1, LEDsupply, Randolph, VT, USA) and 740 nm LEDs (product no. MTE1074N1-R, Marktech Optoelectronics Inc., Latham, NY, USA). All light-sensitive manipulations were done under green safelight.

### Lethality assays of SCRaMbLEd yeast cultures

For lethality assays, cultures were treated as described above in ‘Light-induced recombination experiments’. Hundred µL of each culture were used to prepare sequential dilutions in a 96-well microtiter plate and 20 µL of each dilution was spotted on SD-Leu plates. Assays were performed in three independent replicates. Colonies per spot were counted and converted to colony forming units per microliter according to dilution and drained volume.

### qPCRTag generation

Diagnostic qPCR primer pairs for synIII were designed using a prototype of qTagGer and manually curated to remove redundancies. qTagGer later evolved into a command line tool for *S. cerevisiae* written in python which could be expanded to other organisms. Briefly qTagGer integrates Primer3 for primer design (Koressaar and Remm, 2007; Untergasser *et al*., 2012) with a masking function for *S. cerevisiae* (Koressaar *et al*., 2018) to exclude error prone regions before primer design is performed. Off-target detection is performed by applying bowtie (Langmead *et al*., 2009) to detect all sequences with multiple matches in the genome with a defined mismatch value (default value ≤ 4 bp). All off-target and Primer3 config parameters can be customized in the corresponding .yaml file. qTagGer was subsequently used to predicted loxTags and roxTags for all synthetic chromosomes; references used are listed in Table S6. LoxTags and roxTags were manually curated and are provided in the supplementary data (Supporting Data S1). qTagGer is maintained and available at GitHub (https://github.com/RGSchindler/qTagGer).

### qPCR tagging and data analysis

qPCR analysis of both LoxTags and PCRTags was done by adapting an earlier established protocol (Mitchell *et al*., 2015). Briefly, dispensing of LoxTags and PCRTags was performed using an Echo525 (Labcyte). 50 nL of 100 µM pre-mixed forward and reverse primer were dispensed in 384 PCR plates (Sarstedt, 72.1984.202). Primer plates were prepared in bulk and stored up to four weeks. To perform the qPCR assay 0.95 µL qPCR mastermix (M3003, NEB) containing 2 ng/µL template DNA was dispensed using a nanoliter dispenser (Cobra 4-channel dispenser, ARI). Plates were spun down 500 *g* for 5 min followed by sealing with optical clear permanent seal (Agilent, 24212-001) using the Plateloc Thermal Microplate Sealer (Agilent). qPCRs were run using the Applied Biosystems QuantStudio 5. Samples were pre-incubated at 50 °C (1.6 °C/s) for 2 min followed by 1 min 95 °C (2.57 °C/s). Two-step PCR stage was performed with 30 cycles each 95 °C (2.57 °C/s) 1 s, 67 °C (2 °C/s) 1 min, with single acquisition. Melting curve was acquired from 97 °C (0.1 °C/s) with continuous acquisition. Raw data was exported and analyzed by using a custom R script applying ggPlot2 for heatmap generation.

### *in silico* qPCR tagging

To visualize loxTag pattern of assembled genomes we generated the custom script visLoxP using python. VisLoxP creates *in silico* amplicons based on an input query sequence in fasta format and a .csv file of the oligonucleotide pairs. VisLoxP tests if a respective amplicon can be generated with user defined requirements for the size of the amplicon and the number of accepted primer mismatches (due to not perfect long-read assemblies). If a primer pair matches the defined requirements the value 1 is assigned if not 0. The program gives out a text file which can be further used for visualization. In this study we used an amplicon size of ≤ 1000 bp and allowed ≤ 2 mismatches per primer. The resulting values were subsequently plotted as a heatmap using seaborn (Waskom, 2021). VisLoxP is available at GitHub (https://github.com/RGSchindler/visLoxP).

### High molecular weight DNA extraction and whole genome Nanopore sequencing

Genomic DNA was obtained using the NucleoBond HMW DNA kit (740160.20, Macherey-Nagel, Düren, Germany) according to the manufacturer guidelines using lyticase (#L4025, Sigma) for cell lysis (25 µL; 10,000 U/mL) for 1 h at 37 °C in 1.5 mL Y1 buffer (1 M Sorbitol, 100 mM EDTA pH 8.0, 14 mM β-mercaptoethanol. DNA quality and concentration were assessed via NanoDrop™ 8000 Spectrophotometer and Qubit 3 Fluorometer using dsDNA BR reagents. Library preparation was performed using the Ligation Sequencing kit SQK-LSK109 (Oxford Nanopore Technologies) with the Native Barcoding kit EXP-NDB104 and EXP-NDB114 for multiplexing. Kits were used according to the manufacturers’ guidelines, except the input DNA was increased 5-fold to match the molarity expected in the protocol as no DNA shearing was applied. Sequencing was performed on a MinION Mk1B device using MinION Flow Cells [FLO-MIN111 (R10)] or Flongle Cells [FLO-FLG001 (R9.4.1)].

### Nanopore sequencing data analysis

Basecalling was performed using Guppy (version: 6.0.1+652ffd179) and reads were mapped against the *S. cerevisiae* S288c reference (PRJNA128) with chromosome III replaced by synIII (PRJNA351844) using minimap2 (version: 2.17-r941) (Li, 2018). Canu (version: 2.2) (Koren *et al*., 2017) was used to perform *de novo* assembly of linear chromosomes and Flye (version 2.9.1-b1780) (Kolmogorov *et al*., 2019) to assemble circular synIII derivatives. *De novo* assembled synIII contigs were manually curated using the *de novo* assembly as reference to map the raw reads. Subsequent calling of structural variations (SVs) was performed using sniffles (version: 2.0.7) (Sedlazeck *et al*., 2018). Detected SVs were curated manually, the curation steps of mapping and SV calling were performed until no further SVs could be detected. Obtained synIII sequences were used to generate dot plots in comparison to the respective parental synIII using Mummer (version: 3.5) (Delcher *et al*., 1999). SCRaMbLEgrams for visualization purposes were constructed using the python library pygenomeviz from Python (version: 3.9.7). All raw reads are deposited at BioProject PRJNA884617 and an overview of sequenced strains is given in Table S1.

### Pulsed-field gel electrophoresis

Plug preparation and PFGE was performed as described previously (Schindler *et al*., 2022) according to described methods by Hage and Houseley (2013) (Hage and Houseley, 2013) with the following slight alterations. PFGE was undertaken by running samples on a 1.0% agarose gel in 0.5 X TBE solution at 14 °C on a Bio-Rad clamped homogeneous electric field apparatus (CHEF-DR III, BioRad). 6 V/cm were used with 15 h switch time of 60 s followed by 15 h 120 s at 120°. The resulting gel was stained with 1 X SYBR Safe (ThermoFisher Scientific) and imaged using Typhoon RGB laser scanning system. The known karyotype of BY4741 served as a size standard.

### Flow cytometry

Ploidy of yeast cells was determined using SYTOX Green stained fixed cells in a BD Fortessa Flow Cytometer. Briefly, cells were grown to an OD_600_ of approx. 0.5 with 2 mL of cells collected (4 min at 2,000 *g*), washed with filtered H_2_O and fixed in 1 mL 70% EtOH. Fixed cells were washed twice in 1 mL filter sterilized 50 mM sodium citrate followed by resuspension in 1 mL 50 mM sodium citrate with RNase A (final conc. 0.25 mg/mL) and incubation at 50 °C for 1 h. Subsequently Proteinase K was added with a final conc. of 0.4 mg/mL followed by 1 h at 50 °C before harvesting the cells (4 min 2,000 *g*). Cell pellets were resuspended in 1 mL 50 mM sodium citrate with Sytox Green (5 mM stock; 1:5,000 diluted) and subsequently measured in the BD Fortessa Flow Cytometer. Known haploid (BY4741) and diploid (BY4743) control strains were used as standards.

### Phenotyping on solid and in liquid media

Selected strains were recovered from cryocultures and streaked on YEPD to obtain single colonies. Individual clones were grown in 5 mL YEPD for 40 h. Cultures were measured and normalized to an OD_600_ = 1. 10-fold dilution series were prepared in 96-well microtiter plates and the Rotor HDA+ (Singer Instruments) was used to perform 7x7 grit pinning on the described solid media. Plates were cultivated at 30 °C over a time course of five days. Images were taken every 24 h using the PhenoBooth+ (Singer Instruments).

## Supporting information

Supporting Data S1 - predicted primer

Supporting Data S1 - Genbank sequences

## Data availability statement

The data underlying this study are available in the published article and its online supplementary material. Sequencing raw reads are available under BioProject PRJNA884617. Custom scripts are available at GitHub as indicated. All material created within this study is available from the corresponding authors upon request.

## Author Contributions

LH and DS conceived, planned and designed the study. TAL, MCSO, KS, AARR, LH and DS performed experiments and analyzed the data. CKB, MCSO and DS generated and applied the computational tools. LH and DS wrote the manuscript with the input of all authors. All authors approved the final manuscript.

## Acknowledgments

We thank the Hochrein and Schindler research groups and the MaxGENESYS biofoundry team for fruitful and inspiring discussions. We thank Silvia González Sierra and Gabriele Malengo of the Flow Cytometry & Imaging Facility at the Max Planck Institute for Terrestrial Microbiology as well as Giovanni Scarinci for support in regard to flow cytometry data generation and analysis. We thank Tania Köbel for technical support throughout the study. Jef Boeke (NYU), Yizhi Cai (UoM), Tom Ellis (Imperial College), Leslie Mitchell (NYU) and Torsten Waldminghaus (TU Darmstadt) for sharing strains and/or plasmids. We thank Sally Jones and Bernd Müller-Röber for critical reading of the manuscript.

## Supplementary Material

- Supporting Data S1: LoxTag prediction for all synthetic chromosomes (S1_Lindeboom_predicted_primer.xls).
- Supporting Data S2: Annotated .gb files of all constructed plasmids and the circular synIII (S2_Lindeboom_genbank.zip).

## Data Availability Statement

- Nanopore Sequencing data is available at PRJNA884617.
- Code for primer designer is available at https://github.com/RGSchindler/qTagGer.
- Code for loxTag visualizer is available at https://github.com/RGSchindler/visLoxP.
- L-SCRaMbLE plasmids are available at Addgene, ID #100539 and #100537.
- All within this study created materials are available from the corresponding authors upon request.

## Supporting information

**Figure S1.**
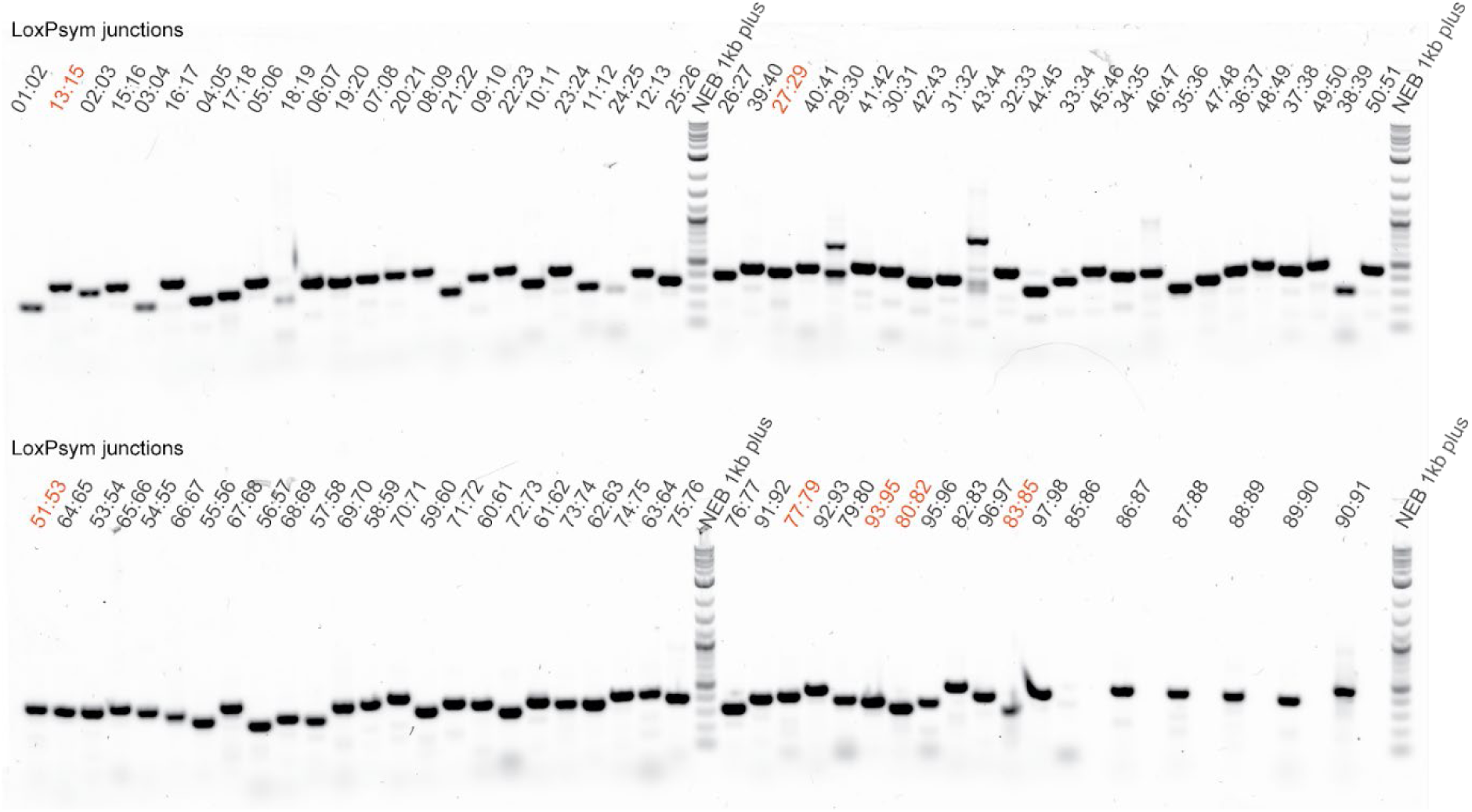
LoxTag amplicon test with standard Taq polymerase. Initial test of the loxTags with standard Taq polymerase (OneTaq 2X Master Mix, NEB) and synIII gDNA as template. For maximal resolution and sensitivity the Typhoon RGB laser scanning system was used to identify unspecific binding. Notably no negative control can be performed because the primer pairs bind in both the synthetic and wild type chromosome III. Gel scan shows individual bands of the loxTags with the expected sizes (*cf*. Table S5). Significant off-target amplicons are only visible for *loxPsym* junction [29:30] and [43:44]. These initial results were later used to optimize the qPCR conditions resulting in the 1 µL total volume protocol described in the material and methods section. Amplicons spanning 2 *loxPsym* sites are indicated in orange.

**Figure S2.**
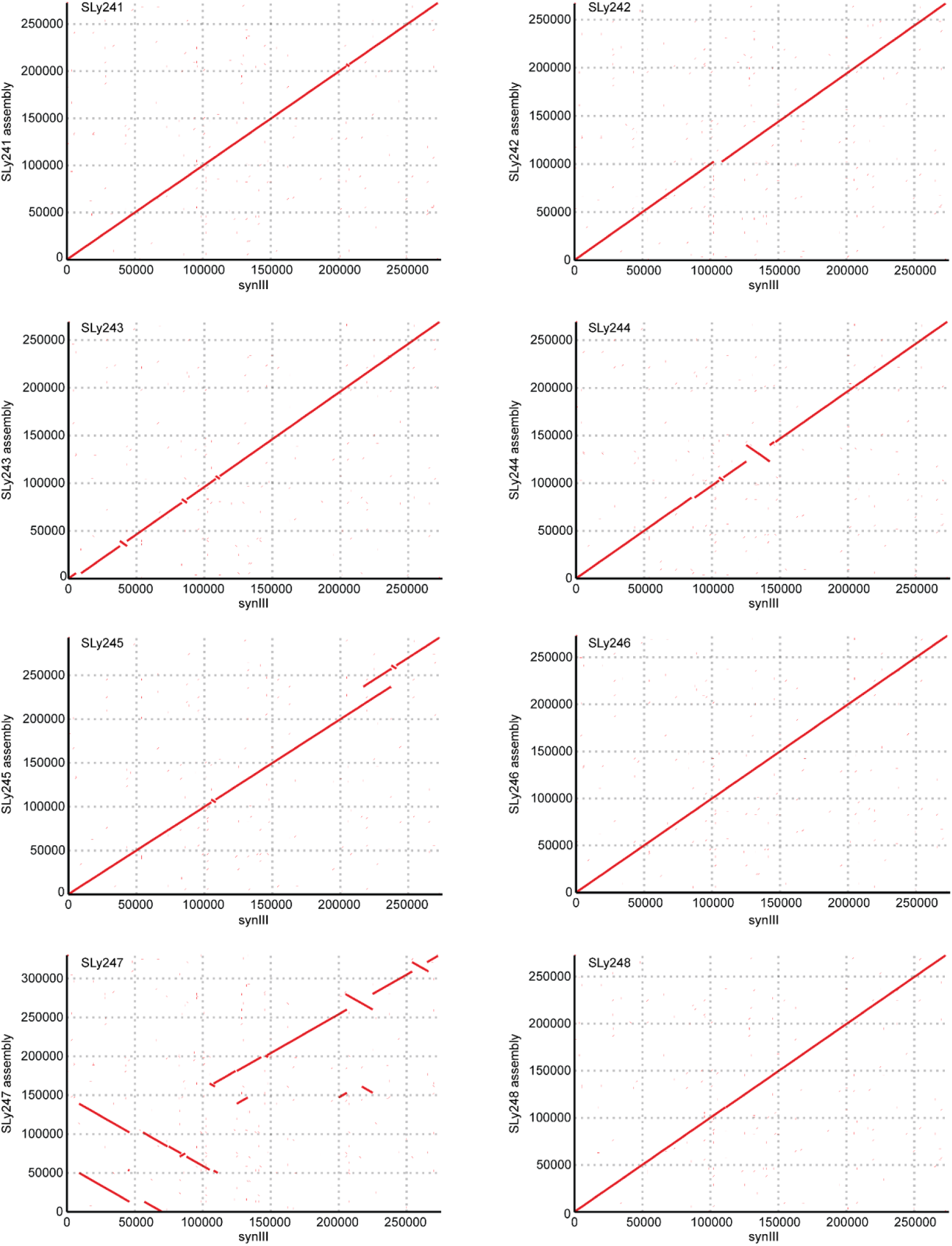
Dot plots of Y1511 synIII derivatives in comparison to the reference synIII. The dot plots visualize structural variations of the SCRaMbLEd isolates in comparison to the synIII reference strain. X-axis corresponds to the genomic positions of the synIII reference in bp and y-axis corresponds to the indicated SCRaMbLEd isolate. SLy247 shows the most complex structural rearrangement including all four types of SCRaMbLE events.

**Figure S3.**
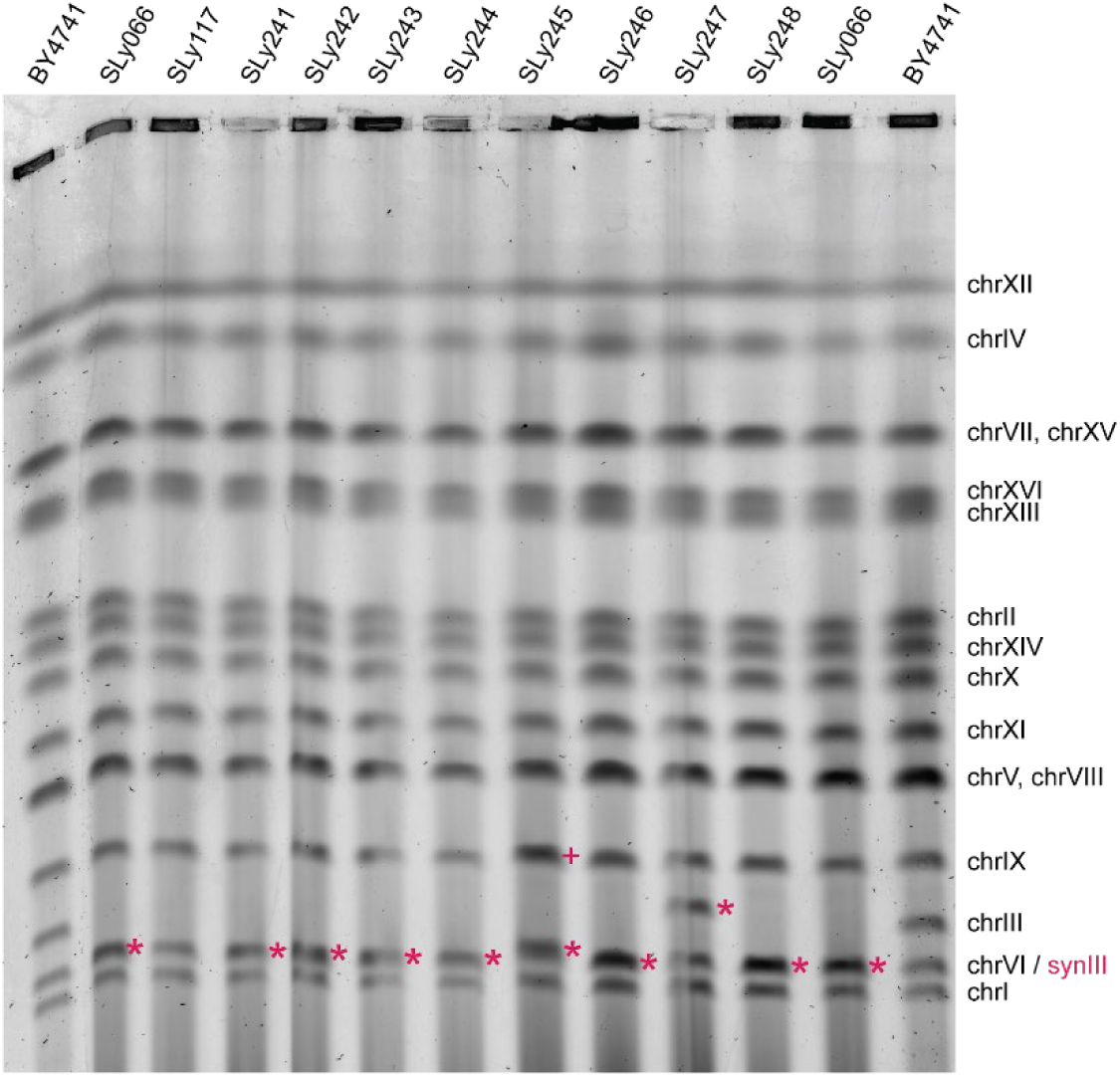
Full image of pulsed-field gel electrophoresis of SCRaMbLEd linear synIII strains and respective controls (*cf*. Fig. 4). Besides the alteration of synIII an increased intensity for chrIX in strain SLy245 is present which matches the Nanopore sequencing data in regard to an aneuploidy for chrIX (*cf*. Fig. 6). * indicates synIII, + indicates chrIX aneuploidy.

**Figure S4.**
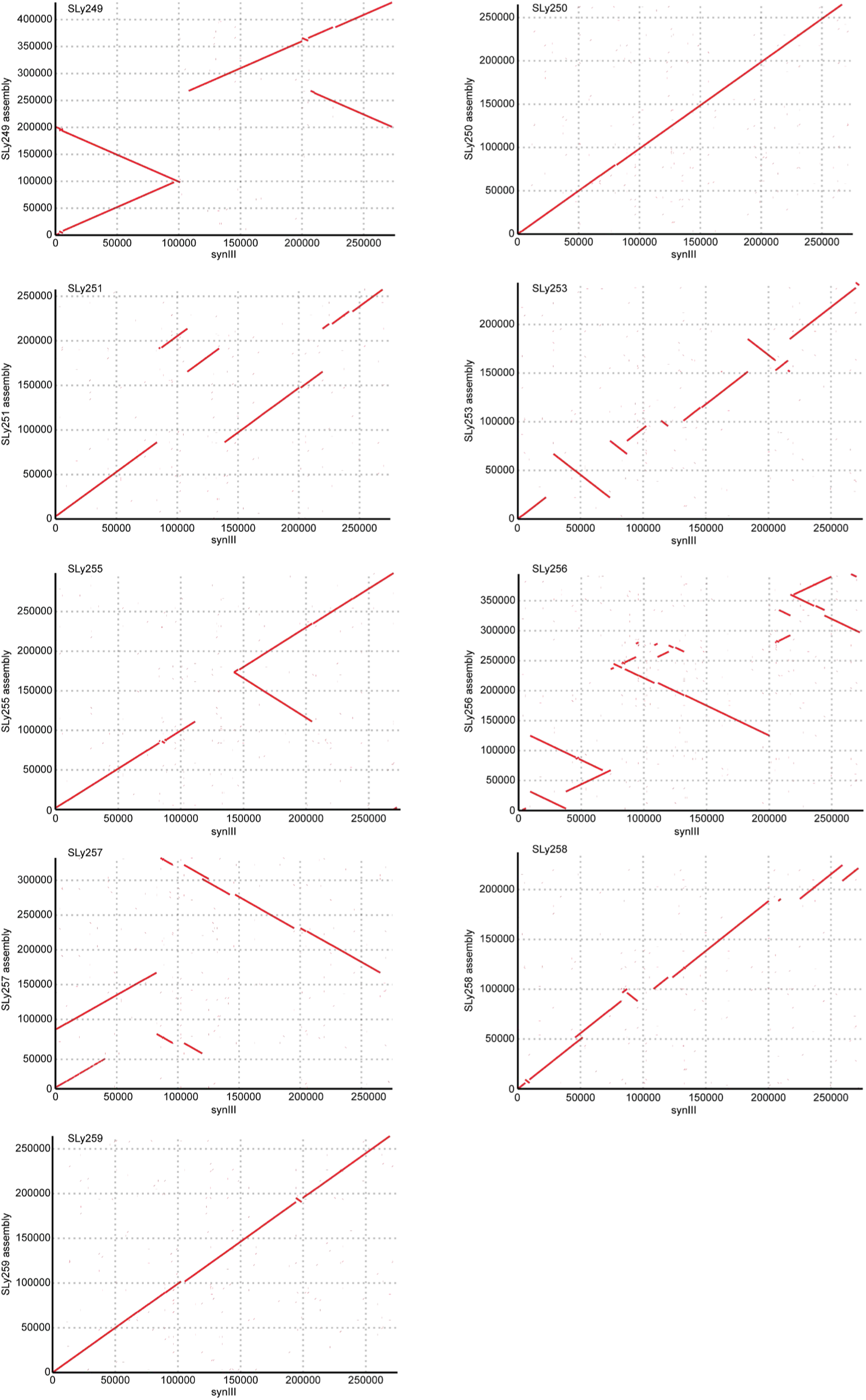
Dot plots of Y1512 derivatives in comparison to the parental strain. The dot plots visualize structural variations of the SCRaMbLEd isolates in comparison to the synIII reference strain. X-axis corresponds to the genomic positions of the synIII reference in bp and y-axis corresponds to the indicated SCRaMbLEd isolate.

**Figure S5.**
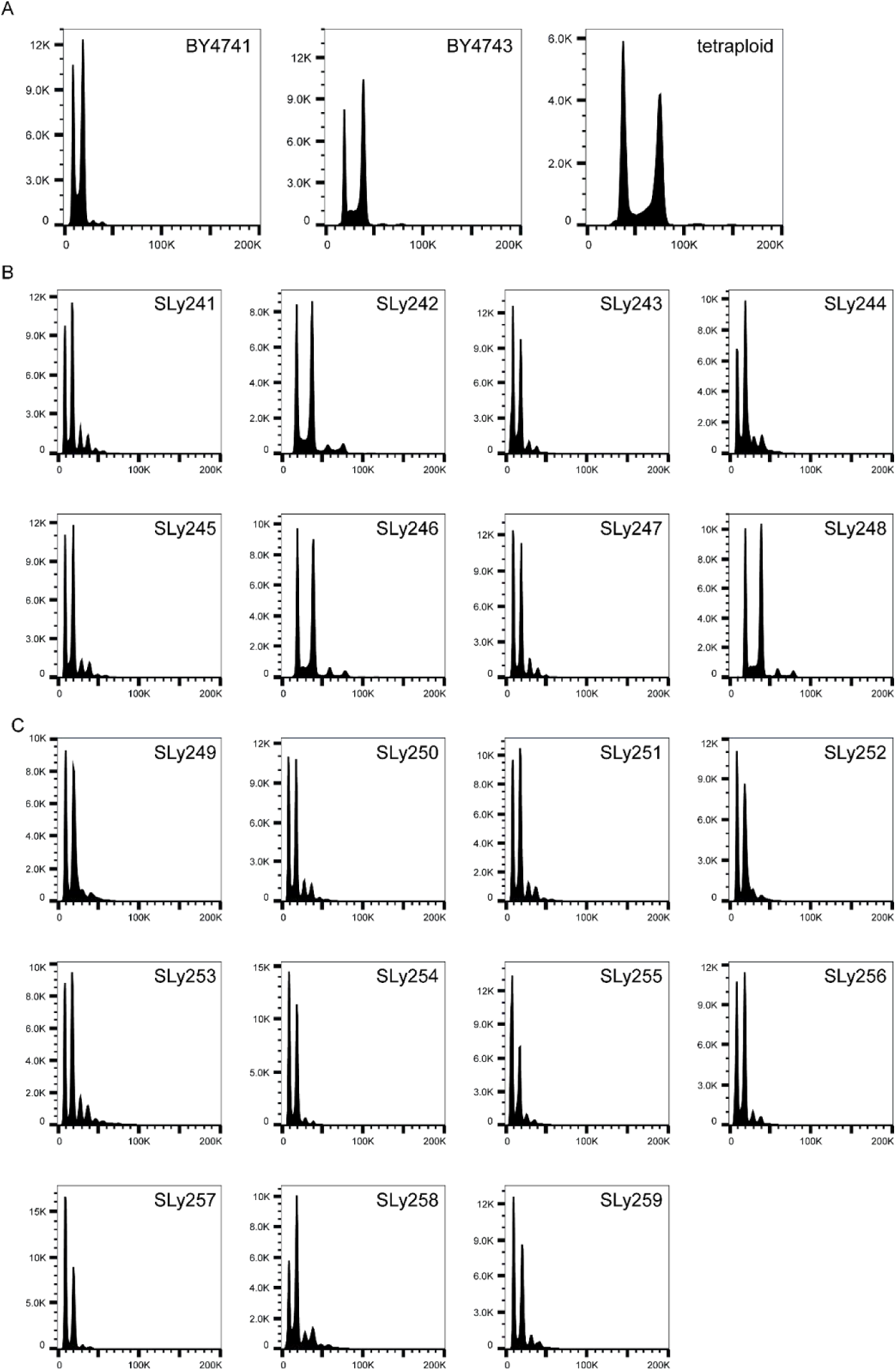
Ploidy analysis of all sequenced isolates in comparison to reference strains BY4741, BY4743 and a tetraploid strain. **(A)** Control strains visualizing known ploidies and serving as standard for the flow cytometry based DNA content determination. Haploid (BY4741), diploid (BY4743) and tetraploid (YCy2990; Schindler & Cai unpublished). **(B)** SLy242, SLy246 and SLy248 of the linear synIII derivatives show a diploid DNA content after SCRaMbLE and indicate a whole genome duplication (no aneuploidies are detected in the NGS data). All remaining strains have a haploid genotype except for SLy245 where an aneuploidy was detected by Nanopore sequencing (*cf*. Fig. 6). **(C)** None of the circular synIII derivatives indicate a whole genome duplication, all strains are haploid. However, a partial chromosome duplication (SLy251) and aneuploidies were detected by Nanopore sequencing (SLy255 and SLy257; *cf*. Fig. 6).

**Tab. S1.** Details of sequenced strains all data are deposited at PRJNA884617.

| Strain ID | Genome coverage* | Average read length | Max. read length | SRA accession |
| --- | --- | --- | --- | --- |
| SLy241 | 44.17 | 17,027 | 319,665 | SAMN32112756 |
| SLy242 | 16.36 | 18,705 | 182,489 | SAMN32112757 |
| SLy243 | 39.75 | 19,254 | 210,066 | SAMN32112758 |
| SLy244 | 42.44 | 15,830 | 200,396 | SAMN32112759 |
| SLy245 | 33.77 | 20,714 | 215,503 | SAMN32112760 |
| SLy246 | 38.84 | 25,051 | 253,179 | SAMN32112761 |
| SLy247 | 50.05 | 15,783 | 226,520 | SAMN32112762 |
| SLy248 | 73.37 | 19,836 | 227,007 | SAMN32112763 |
| SLy249 | 36.35 | 18,346 | 215,975 | SAMN32112764 |
| SLy250 | 16.17 | 17,867 | 248,777 | SAMN32112765 |
| SLy251 | 26.14 | 14,486 | 230,528 | SAMN32112766 |
| SLy252 | 17.05 | 22,474 | 167,415 | SAMN32112767 |
| SLy253 | 18.4 | 21,260 | 209,689 | SAMN32112768 |
| SLy254 | 29.26 | 20,652 | 253,720 | SAMN32112769 |
| SLy255 | 34.77 | 28,742 | 278,058 | SAMN32112770 |
| SLy256 | 21.82 | 18,307 | 232,287 | SAMN32112771 |
| SLy257 | 56.38 | 25,174 | 291,363 | SAMN32112772 |
| SLy258 | 51.09 | 15,073 | 164,359 | SAMN32112773 |
| SLy259 | 14.65 | 12,089 | 227,073 | SAMN32112774 |

\* based on the average coverage of chromosome 1 to 16, excluding mtDNA.

**Tab. S2.** Yeast strains generated and used in this study.

| Yeast strain | Genotype [plasmid] | Parental strain | Reference |
| --- | --- | --- | --- |
| BY4741 | MATa leu2Δ0 met15Δ0 ura3Δ0 his3Δ1 | NA | (Brachmann *et al.*, 1998) |
| BY4743 | MATa/α his3Δ1/his3Δ1 leu2Δ0/leu2Δ0 LYS2/lys2Δ0 met15Δ0/MET15 ura3Δ0/ura3Δ0 | NA | (Brachmann *et al.*, 1998) |
| synIII | MATα his3Δ1 leu2Δ0 lys2Δ0 ura3Δ0 synIII ho::syn.SUP61-URA3 | NA | (Annaluru *et al.*, 2014) |
| SLy053 | MATα his3Δ1 leu2Δ0 lys2Δ0 ΔURA3 synIII ho::syn.SUP61-URA3 | synIII | this study |
| SLy066 | MATα his3Δ1 leu2Δ0 lys2Δ0 ΔURA3 synIII ho::syn.SUP61-ΔURA3 | SLy053 | this study |
| SLy117 | MATα his3Δ1 leu2Δ0 lys2Δ0 ΔURA3 synIII ring chromosome ho::syn.SUP61-ΔURA3 | SLy066 | this study |
| Y1426 | [pLH_Scr15 + pLM494] | SLy066 | this study |
| Y1510 | [pLH_Scr15] | BY4742 | this study |
| Y1511 | [pLH_Scr15] | SLy066 | this study |
| Y1512 | [pLH_Scr15] | SLy117 | this study |
| SLy241 | SLy066 SCRaMbLEd | Y1511 | this study |
| SLy242 | SLy066 SCRaMbLEd | Y1511 | this study |
| SLy243 | SLy066 SCRaMbLEd | Y1511 | this study |
| SLy244 | SLy066 SCRaMbLEd | Y1511 | this study |
| SLy245 | SLy066 SCRaMbLEd | Y1511 | this study |
| SLy246 | SLy066 SCRaMbLEd | Y1511 | this study |
| SLy247 | SLy066 SCRaMbLEd | Y1511 | this study |
| SLy248 | SLy066 SCRaMbLEd | Y1511 | this study |
| SLy249 | SLy117 SCRaMbLEd | Y1512 | this study |
| SLy250 | SLy117 SCRaMbLEd | Y1512 | this study |
| SLy251 | SLy117 SCRaMbLEd | Y1512 | this study |
| SLy252 | SLy117 SCRaMbLEd | Y1512 | this study |
| SLy253 | SLy117 SCRaMbLEd | Y1512 | this study |
| SLy254 | SLy117 SCRaMbLEd | Y1512 | this study |
| SLy255 | SLy117 SCRaMbLEd | Y1512 | this study |

| **Yeast strain** | **Genotype [plasmid]** | **Parental strain** | **Reference** |
| --- | --- | --- | --- |
| SLy256 | SLy117 SCRaMbLEd | Y1512 | this study |
| SLy257 | SLy117 SCRaMbLEd | Y1512 | this study |
| SLy258 | SLy117 SCRaMbLEd | Y1512 | this study |
| SLy259 | SLy117 SCRaMbLEd | Y1512 | this study |
| SLy273 | SLy066 SCRaMbLEd | Y1426 | this study |
| SLy274 | SLy066 SCRaMbLEd | Y1426 | this study |
| SLy275 | SLy066 SCRaMbLEd | Y1426 | this study |

**Tab. S3.**
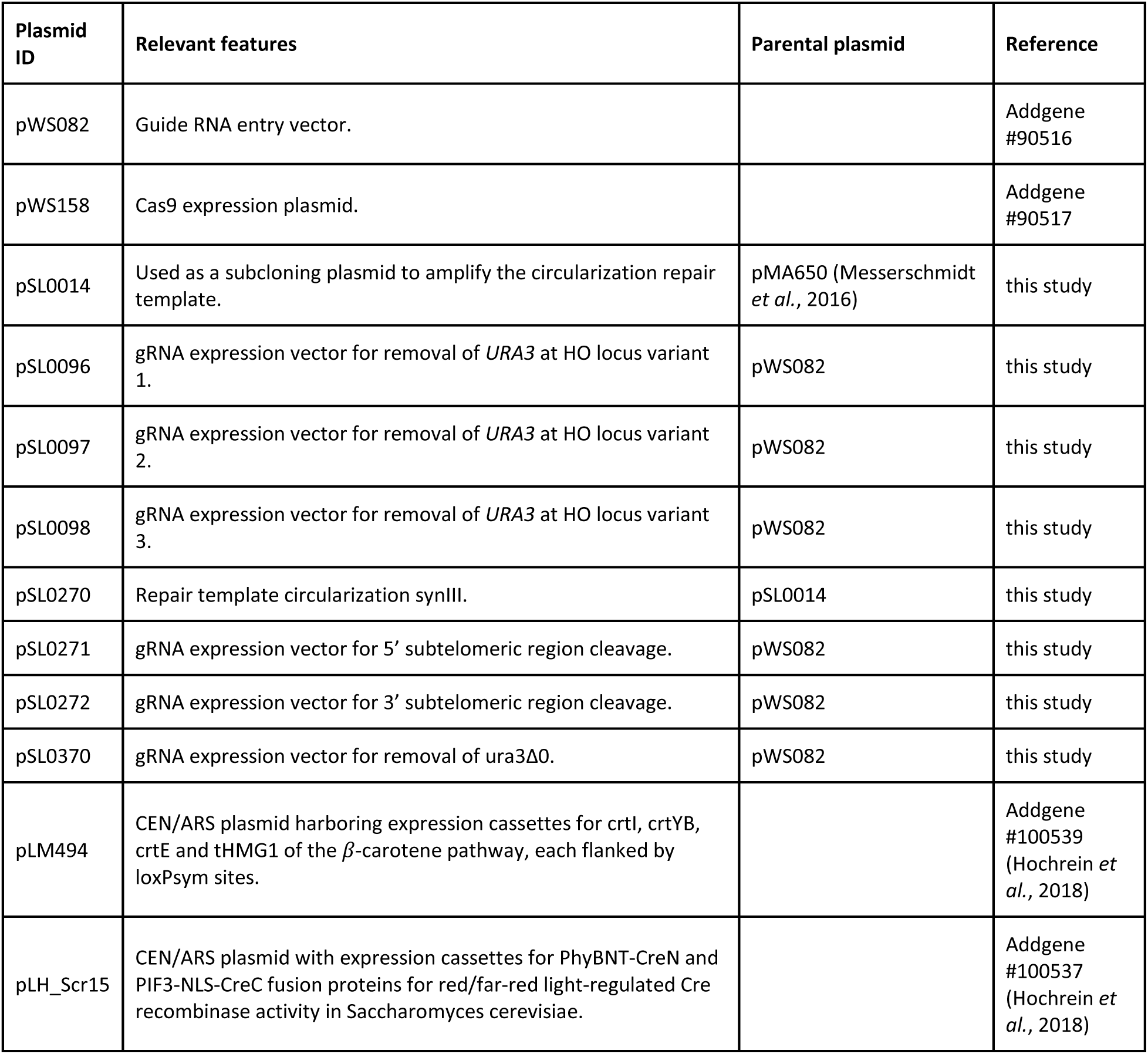
Plasmids constructed and used in this used in this study.

| Plasmid ID | Relevant features | Parental plasmid | Reference |
| --- | --- | --- | --- |
| pWS082 | Guide RNA entry vector. |  | Addgene #90516 |
| pWS158 | Cas9 expression plasmid. |  | Addgene #90517 |
| pSL0014 | Used as a subcloning plasmid to amplify the circularization repair template. | pMA650 (Messerschmidt *et al.*, 2016) | this study |
| pSL0096 | gRNA expression vector for removal of *URA3* at HO locus variant 1. | pWS082 | this study |
| pSL0097 | gRNA expression vector for removal of *URA3* at HO locus variant 2. | pWS082 | this study |
| pSL0098 | gRNA expression vector for removal of *URA3* at HO locus variant 3. | pWS082 | this study |
| pSL0270 | Repair template circularization synIII. | pSL0014 | this study |
| pSL0271 | gRNA expression vector for 5' subtelomeric region cleavage. | pWS082 | this study |
| pSL0272 | gRNA expression vector for 3' subtelomeric region cleavage. | pWS082 | this study |
| pSL0370 | gRNA expression vector for removal of *ura3Δ0*. | pWS082 | this study |
| pLM494 | CEN/ARS plasmid harboring expression cassettes for crtI, crtYB, crtE and tHMG1 of the β-carotene pathway, each flanked by loxPsym sites. | | Addgene #100539 (Hochrein *et al.*, 2018) |
| pLH_Scr15 | CEN/ARS plasmid with expression cassettes for PhyBNT-CreN and PIF3-NLS-CreC fusion proteins for red/far-red light-regulated Cre recombinase activity in *Saccharomyces cerevisiae*. |  | Addgene #100537 (Hochrein *et al.*, 2018) |

**Tab. S4.**
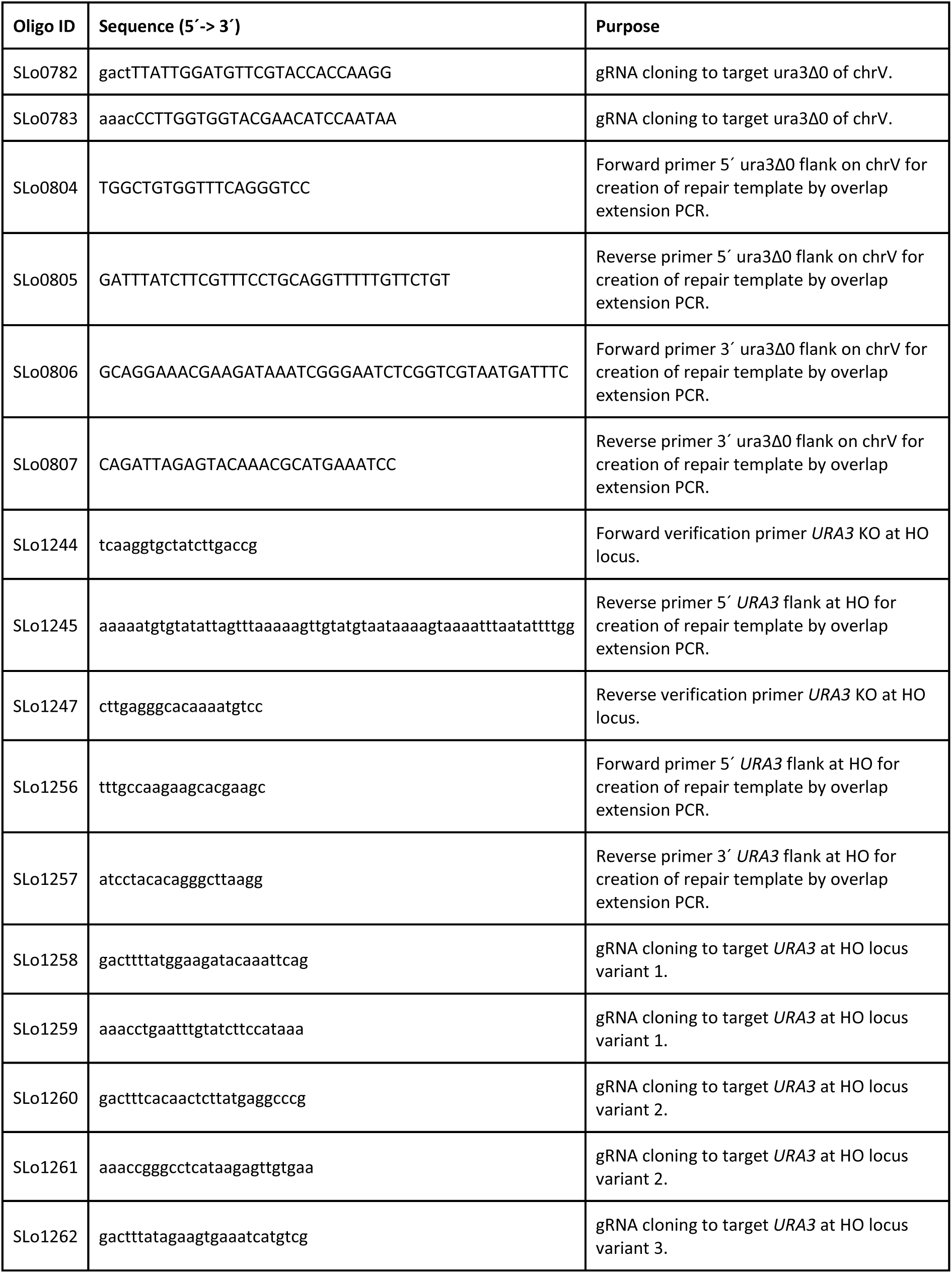
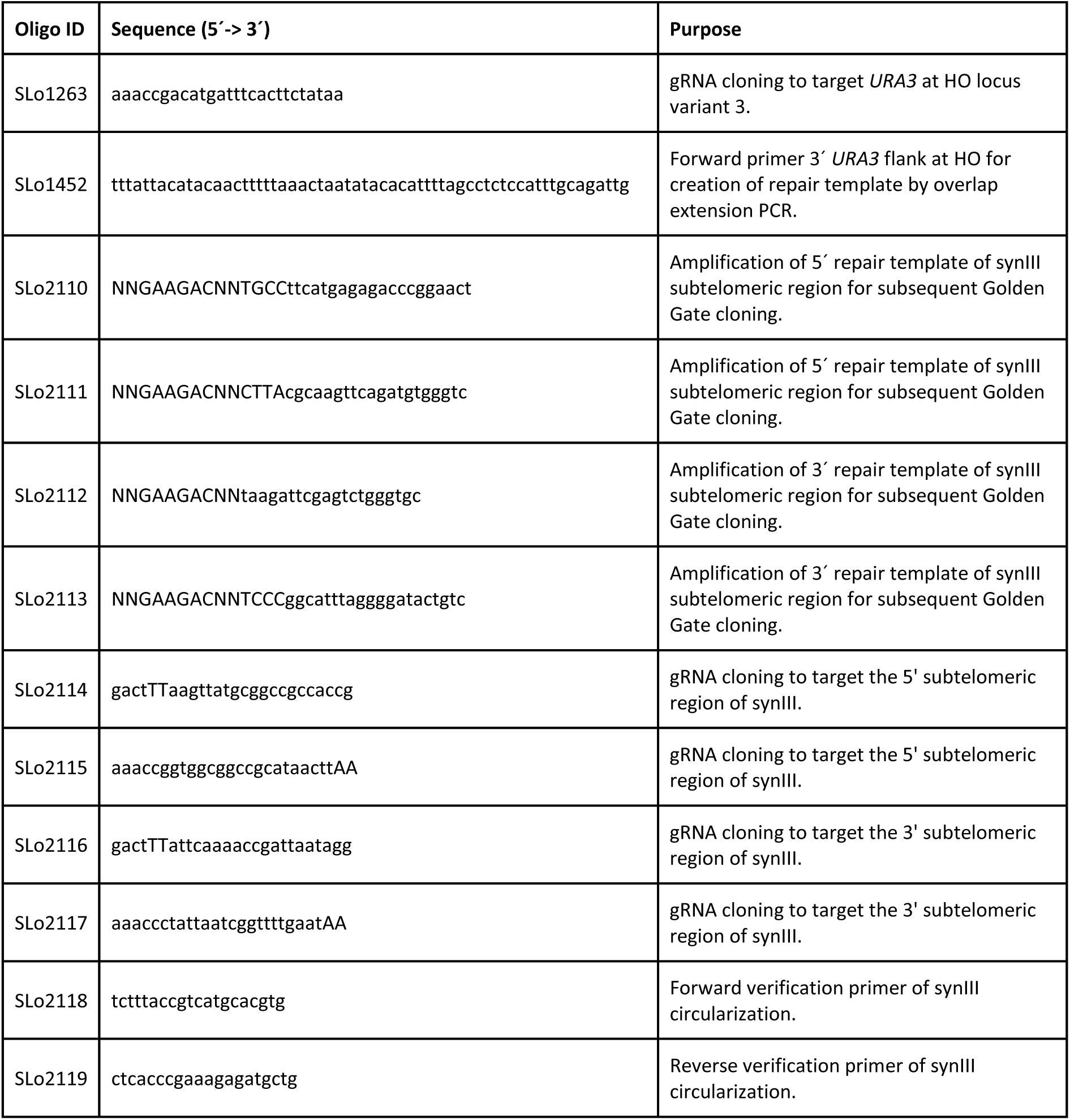
Oligonucleotides used in this study.

**Tab. S5.**
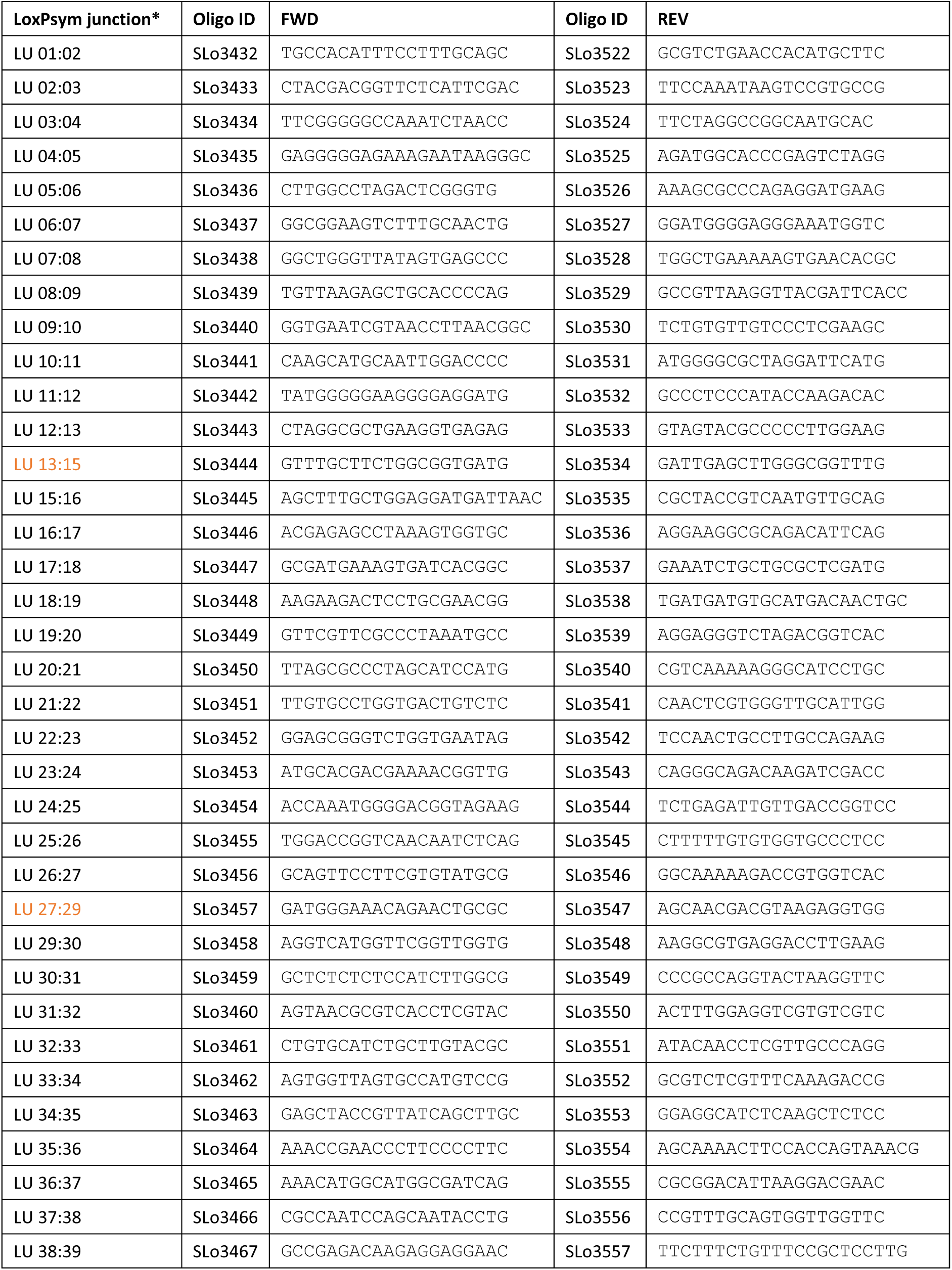
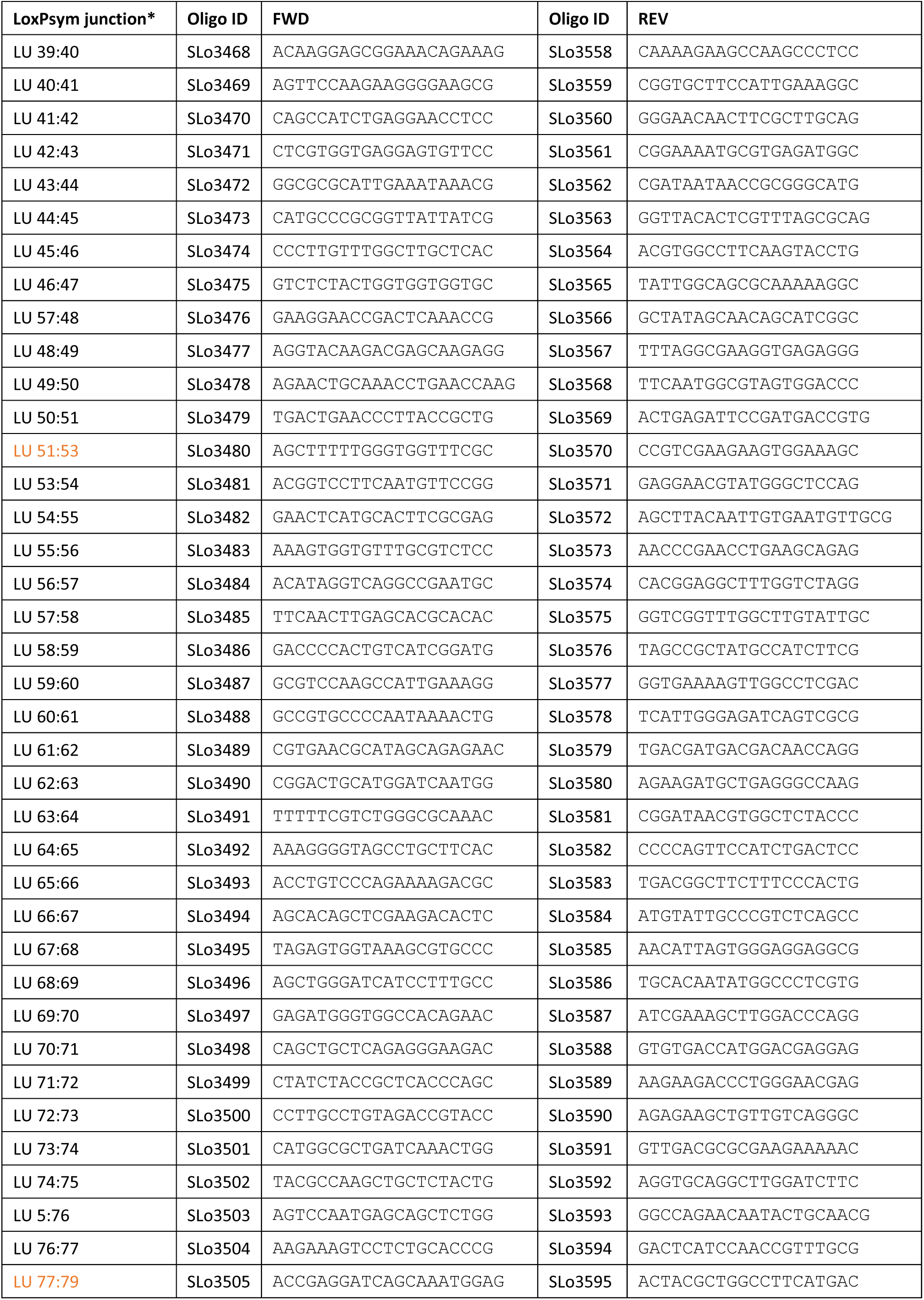
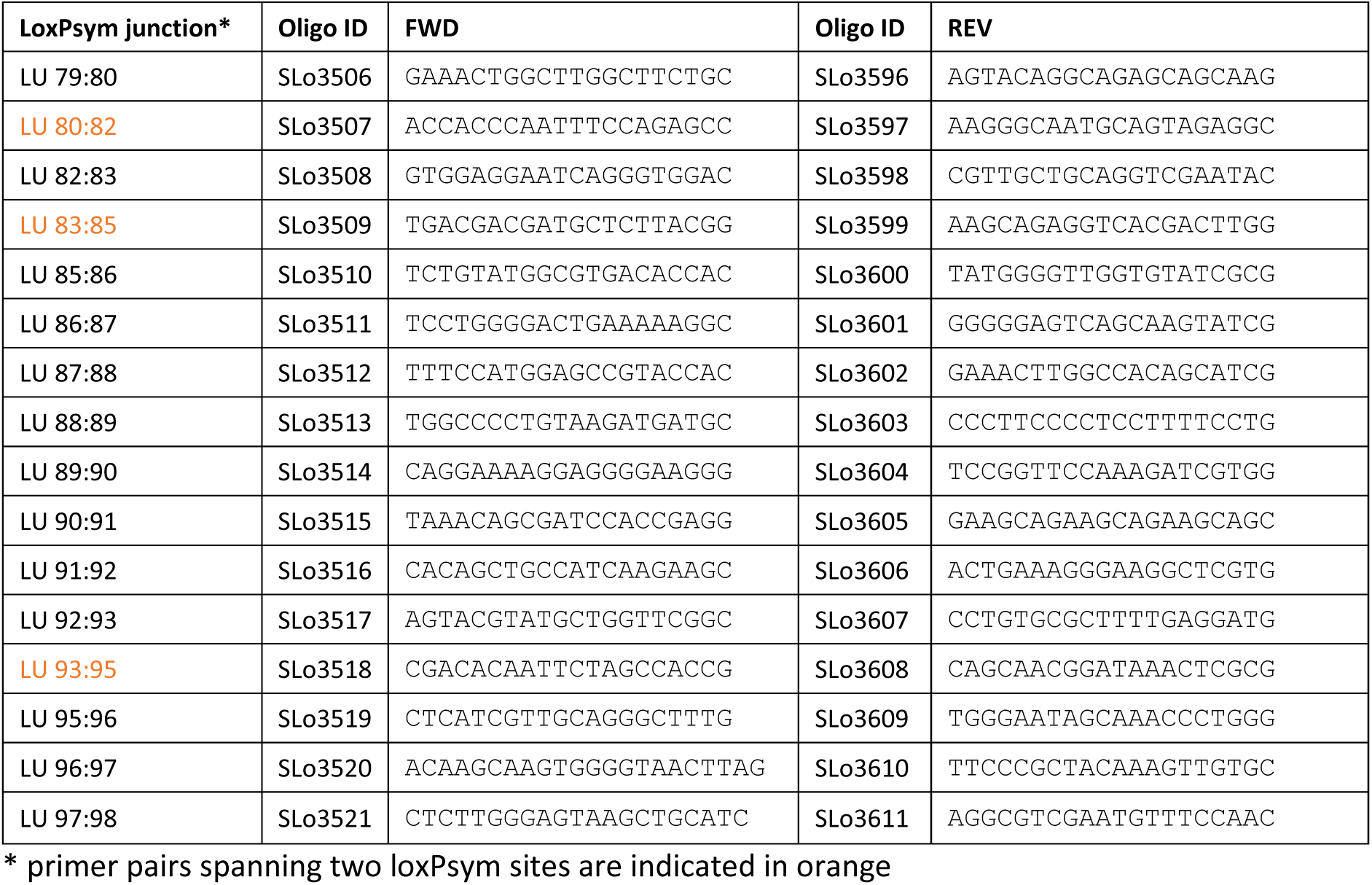
**synIII loxTags used in this study.** All other loxTags and the roxTags are provided with the supporting data (Supporting Data S1).

**Tab. S6.** References used for loxTag generation.

| **Chromosome** | **Reference** |
| --- | --- |
| synI | chr01_3_24 |
| synII | chr02_3_26 |
| synIII | chr03_9_02 |
| synIV | chr04_3_71 |
| synV | chr05_3_43 |
| synVI | chr06_3_27 |
| synVII | chr07_3_60 |
| synVIII | chr08_3_35 |
| synIX | chr09_3_54 |
| synX | chr10_9_01 |
| synXI | chr11_3_38 |
| synXII | chr12_9_05 |
| synXIII | chr13_3_43 |
| synXIV | chr14_3_29 |
| synXV | chr15_3_45 |
| synXVI | chr16_3_43 |
| tRNA neochromosome | PRJNA351844 |

